# Global nuclear reorganization during heterochromatin replication in the giant-genome plant *Nigella damascena* L.

**DOI:** 10.1101/2023.08.15.552960

**Authors:** Eugene A. Arifulin, Dmitry V. Sorokin, Nadezhda A. Anoshina, Maria A. Kuznetsova, Anna A. Valyaeva, Daria M. Potashnikova, Denis O. Omelchenko, Veit Schubert, Tatyana D. Kolesnikova, Eugene V. Sheval

**Affiliations:** Belozersky Institute of Physico-Chemical Biology, Lomonosov Moscow State University, Moscow, Russia; Laboratory of Mathematical Methods of Image Processing, Faculty of Computational Mathematics and Cybernetics, Lomonosov Moscow State University, Moscow, Russia; School of Biomedical Sciences and Pharmacy, University of Newcastle, NSW, Australia; School of Bioengineering and Bioinformatics, Lomonosov Moscow State University, Moscow, Russia; Department of Cell Biology and Histology, School of Biology, Lomonosov Moscow State University, Moscow, Russia; Institute for Information Transmission Problems of the Russian Academy of Sciences, Moscow, Russia; Leibniz Institute of Plant Genetics and Crop Plant Research (IPK) Gatersleben, D-06466 Seeland, Germany; Institute of Molecular and Cellular Biology SB RAS, 630090 Novosibirsk, Russia

**Keywords:** genome size, chromatin, chromonema, replication, super-resolution microscopy, *Nigella damascena* L., transmission electron microscopy

## Abstract

Among flowering plants, genome size varies remarkably, by >2200-fold, and this variation depends on the loss and gain of non-coding DNA sequences that form distinct heterochromatin complexes during interphase. In plants with giant genomes, most chromatin remains condensed during interphase, forming a dense network of heterochromatin threads called interphase chromonemata. Using super-resolution light and electron microscopy, we studied the ultrastructure of chromonemata during and after replication in root meristem nuclei of *Nigella damascena* L. During S-phase, heterochromatin undergoes transient decondensation locally at DNA replication sites. Due to the abundance of heterochromatin, the replication leads to a robust disassembly of the chromonema meshwork and a general reorganization of the nuclear morphology visible even by conventional light microscopy. After replication, heterochromatin recondenses, restoring the chromonema structure. Thus, we show that heterochromatin replication in interphase nuclei of giant-genome plants induces a global nuclear reorganization.

## INTRODUCTION

Genomic DNA in eukaryotes is hierarchically organized and compacted within interphase nuclei by interacting with various cations, proteins, and RNAs to form diverse morphological complexes generally referred to as chromatin. In interphase nuclei, chromatin can be divided into two major fractions - euchromatin and heterochromatin. Heterochromatin includes chromosomal regions containing inactive genes and regions enriched for non-coding sequences (including various transposable elements). These morphological findings correlate well with chromosome conformation capture (Hi-C) data indicating that interphase chromatin in animals is partitioned into active A and repressed B compartments (Lieberman-Aiden *et al*., 2009). A and B compartments have also been detected in interphase nuclei of some plants (Dong *et al*., 2017; Chen *et al*., 2020; Montgomery *et al*., 2020; Zhang *et al*., 2021; Li *et al*., 2022).

The distribution and shape of heterochromatin complexes may vary in cells and tissues. In the majority of mammalian species, small blocks of heterochromatin are located preferentially at the nuclear periphery in contact with the nuclear envelope and around nucleoli (perinucleolar chromatin) (Solovei *et al*., 2016; van Steensel and Belmont, 2017). A unique exception, in which euchromatin and heterochromatin swapped positions, was found in rod photoreceptors of nocturnal mammals (Solovei *et al*., 2009; Falk *et al*., 2019; Feodorova *et al*., 2020). However, data obtained mainly from animals, especially mammals, may not reflect chromatin organization in plants. Indeed, three unusual interphase nuclear patterns, differing in the amount, morphology, and localization of heterochromatin, have been described in plants by transmission electron microscopy (TEM). Heterochromatin in different species can form either (i) chromocenters, (ii) chromomeres, or (iii) chromonemata (Nagl and Fusenig, 1979; Nagl and Bachmann, 1980; Nagl *et al*., 1983).

i. In small-genome plants, the heterochromatin forms dense discrete bodies, called chromocenters (Ramachandran and Narayan, 1985; Ceccarelli *et al*., 1998; Fransz and de Jong, 2011; Schubert *et al*., 2012). This type of nuclear organization was demonstrated and studied in detail, in particular, for the model plant *Arabidopsis thaliana* (0.435 Gbp/1C; Schmuths et al. 2004) where the chromocenters contain the highly methylated, mostly repetitive DNA sequences (Fransz *et al*., 2002).
ii. *Nicotiana tabacum* (5.072 Gbp/1C; (Leitch *et al*., 2008) ((Nagl *et al*., 1983) and *Senecio viscosus* (1.519 Gbp/1C; (Nagl *et al*., 1983)) (Nagl, 1985) nuclei contain numerous discrete complexes of condensed chromatin (chromomeres), which are scattered throughout the whole nucleoplasm.
iii. In some plant species, the nucleoplasm is filled with a meshwork of thick anastomosing fibers of condensed chromatin, which were referred to as interphase chromonemata (Sparvoli *et al*., 1965; Nagl and Fusenig, 1979; Nagl, 1985; Hao *et al*., 1994; Kuznetsova *et al*., 2017; Kuznetsova and Sheval, 2016). It should be noted that the term ‘chromonema’ (plural: ‘chromonemata’) was originally used to name helically coiled chromatin threads forming the sister chromatids of large (>12 Mb) metaphase chromosomes during mitosis and meiosis (Baranetzky, 1880; Shinke, 1930; Manton, 1950; Kubalová *et al*., 2023; Câmara *et al*., 2023). The term was introduced by Vejdovsky (1912) from Greek: chrôma - color and nêma - thread. Later, the term was also applied to designate chromatin fibers in interphase nuclei of some plants (Sparvoli *et al*., 1965). In this article, we use the term ‘chromonema’ in the second meaning (for discussion see also (Kuznetsova and Sheval, 2016). To our knowledge, this type of heterochromatin organization was never described in animals, and the chromonema organization is also only scarcely studied in plants.

Heterochromatin can be structurally reorganized during replication. In general, replication leads to the localized decondensation of heterochromatin domains. For example, a transient heterochromatin decondensation at the sites of processive DNA synthesis was demonstrated in cultured human cells (Chagin *et al*., 2019). Intuitively, the high amount of condensed chromatin in plant nuclei showing a chromonema organization may require a more global reorganization of the nucleus, than typical for animal cells.

Here, we examined the interphase nuclear organization in 32 plant species by TEM and found that condensed chromatin forms thick threads (interphase chromonemata) in all giant-genome (>5 Gbp/1C) plants. Then, we studied the chromatin reorganization during and after replication in root meristem nuclei of *Nigella damascena* L. (Ranunculaceae), showing a giant genome size of 10,584.0 Gbp/1C (Olszewska and Osiecka, 1983). Condensed chromatin in *N. damascena* nuclei forms a typical chromonema meshwork, and about half of the genome is composed of tandem repeats (preferentially LTR retrotransposons). Super-resolution microscopy on semithin sections revealed that replication induces chromonema decondensation, leading to the reorganization of the global nuclear morphology. Via correlative light and electron microscopy (CLEM), we revealed that heterochromatin replication occurs within decondensed chromatin, which recondenses when replication is finished.

## RESULTS

### Nuclear organization in plants depends on genome size

We analyzed the ultrastructural organization of root meristematic cells from 32 plant species. In agreement with published data (Nagl *et al*., 1983; Nagl, 1985), we found three variants of the nuclear organization (Figures 1a-c). (i) In the first variant, most of the nucleoplasmic volume was occupied by diffuse euchromatin, heterochromatin was rare and formed small blocks (chromocenters) preferentially in contact with the nuclear envelope (Figure 1a; Figure S1). (ii) In the second variant, heterochromatin was more abundant and formed irregularly shaped complexes (chromomeres) not only in contact with the nuclear envelope but also throughout the nucleoplasm (Figure 1b; Figure S2). (iii) In the third variant, nuclei were filled with a meshwork of thick (150-300 nm) heterochromatin fibers occupying approximately half of the total volume of the nucleoplasm (Figure 1c; Figure S3). Such heterochromatin fibers have been described in the nuclei of several plant species and were termed ‘chromonemata’ (Sparvoli *et al*., 1965; Nagl and Fusenig, 1979; Kuznetsova and Sheval, 2016). Besides the three described categories, transitional variants occur, which made the classification of some species difficult (Figures S1-S3). Nevertheless, most species could be assigned to one of the three categories.

**Figure 1.**
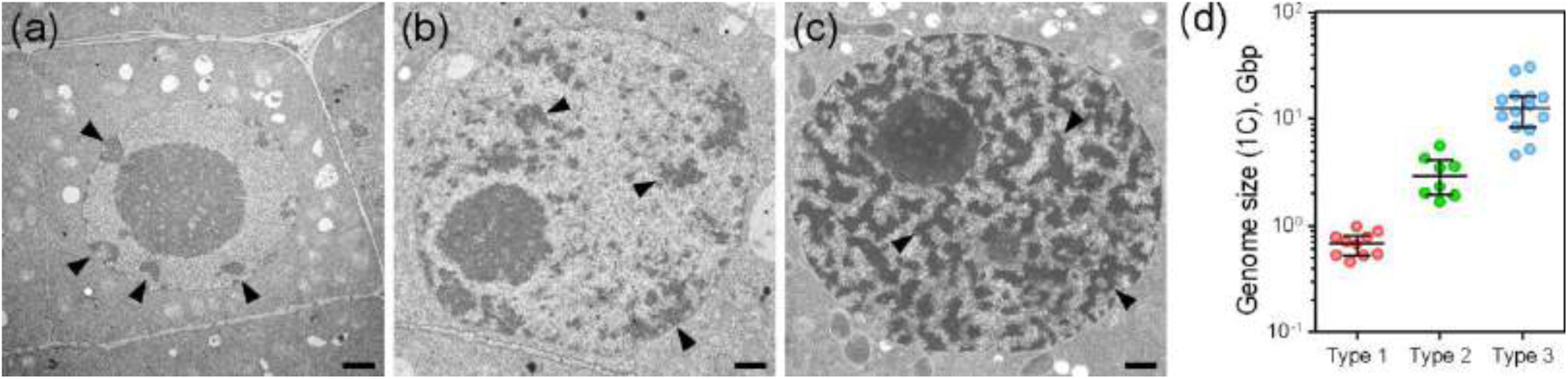
Three variants of heterochromatin organization and distribution in root meristem nuclei of different plants. (a) Nucleus of *Solanum lycopersicum* L. (type 1). The heterochromatin forms only a few discrete complexes (chromocenters) located mainly at the periphery of the nucleus in contact with the nuclear envelope. Heterochromatin is marked with arrowheads. Bars = 1 μm. (b) Nucleus of *Dahlia variabilis* Desf. (type 2). Heterochromatin forms numerous complexes (chromomeres), very heterogeneous in structure and size, localized at the nuclear periphery as well as internally. Heterochromatin is marked with arrowheads. (c) Nucleus of *N. damascena* (type 3). Heterochromatin forms thick branching threads (chromonemata) that fill the entire nucleoplasm. A predominant localization of heterochromatin at the periphery of the nucleus or around the nucleolus, as is typical for animal nuclei, was not observed. Heterochromatin is marked with arrowheads. (d) The dependence of heterochromatin organization on genome size (detailed data are shown in Table 1). Median and interquartile range are shown for each type of heterochromatin distribution.

**Table 1.** Genome size (references from Plant DNA C-values database (https://cvalues.science.kew.org/search)) and heterochromatin organization in plants, evaluated in this study using TEM (see Figures S1, S2, and S3).

| **Species** | **Genome size, per 1C (Gbp)** | **Reference** | **Heterochromatin organization** |
| --- | --- | --- | --- |
| *Daucus carota* L. | 0.461 | (Nowicka *et al.*, 2016) | chromocenters |
| *Brassica rapa* L. | 0.528 | (Johnston *et al.*, 2005) | chromocenters |
| *Cucurbita pepo* L. | 0.529 | (Arumuganathan and Earle, 1991) | chromocenters |
| *Linum usitatissimum* L. | 0.544 | (Marie and Brown, 1993) | chromocenters |
| *Brassica oleracea* L. | 0.671 | (Arumuganathan and Earle, 1991) | chromocenters |
| *Phaseolus vulgaris* L. | 0.701 | (Marie and Brown, 1993) | chromocenters |
| *Catharanthus roseus* (L.) G. Don | 0.784 | (Zonneveld *et al.*, 2005) | chromocenters |
| *Lupinus polyphyllus* Lindl. | 0.784 | (Kubešová *et al.*, 2010) | chromocenters |
| *Cucumis sativus* L. | 0.882 | (Ramachandran and Narayan, 1985) | chromocenters |
| *Solanum lycopersicum* L. | 0.980 | (Obermayer *et al.*, 2002) | chromocenters |
| *Gossypium herbaceum* L. | 1.671 | (Hendrix and Stewart, 2005) | chromomeres |
| *Vicia sativa* L. | 1.911 | (Zonneveld *et al.*, 2005) | chromomeres |
| *Dahlia variabilis* Desf. | 2.024 | (Temsch *et al.*, 2008) | chromomeres |
| *Zea mays* L. | 2.328 | (Arumuganathan and Earle, 1991) | chromomeres |
| *Rudbeckia hirta* L. | 3.479 | (Bai *et al.*, 2012) | chromomeres |
| *Helianthus annuus* L. | 3.597 | (Sanchez *et al.*, 2014) | chromomeres |
| *Lathyrus oleraceus* Lam. | 4.263 | (Zonneveld *et al.*, 2005) | chromomeres |
| *Vicia cracca* L. | 5.586 | (Grime and Mowforth, 1982) | chromomeres |
| *Vicia sepium* L. | 4.606 | (Rees *et al.*, 1966) | chromonemata |
| *Hordeum vulgare* L. | 5.189 | (Jakob *et al.*, 2004) | chromonemata |
| *Lathyrus odoratus* L. | 8.036 | (Murray *et al.*, 1992) | chromonemata |
| *Lathyrus latifolius* L. | 8.389 | (Pustahija *et al.*, 2013) | chromonemata |
| *Nigella sativa* L. | 10.388 | (Bennett and Smith, 1976) | chromonemata |
| *Nigella damascena* L. | 10.584 | (Olszewska and Osiecka, 1983) | chromonemata |
| *Allium fistulosum* L. | 11.642 | (Baranyi and Greilhuber, 1999) | chromonemata |
| *Vicia faba* L. | 13.348 | (Kron and Husband, 2012) | chromonemata |
| *Allium altaicum* Pall. | 15.190 | Ohri D, Fritsch R, Hanelt P., pers. comm. 1996* | chromonemata |
| *Allium sativum* L. | 15.876 | Ohri D, Fritsch R, Hanelt P., pers. comm. 1996* | chromonemata |
| *Allium schoenoprasum* L. | 16.023 | (Zonneveld *et al.*, 2005) | chromonemata |
| *Allium cepa* L. | 16.508 | (Barow and Meister, 2003) | chromonemata |
| *Allium porrum* L. | 28.371 | (Zonneveld *et al.*, 2005) | chromonemata |
| *Allium ramosum* L. | 30.772 | (Ohri *et al.*, 1998) | chromonemata |

\* Cited from Plant DNA C-values database (https://cvalues.science.kew.org/search)

By analyzing the genome size of selected plants from the Plant DNA C-values Database, we found a strong correlation between genome size and nuclear organization (Table 1; Figure 1d). Plants with variant 1 nuclei had relatively small genome sizes (<1.5 Gbp/1C), plants with variant 2 nuclei had large genome sizes (∼1.5-5 Gbps/1C), and plants with chromonemata (variant 3) had giant genome sizes (>5 Gbps/1C). Thus, chromatin organization in plants depends on genome size, and chromonema organization of heterochromatin may be a consequence of giant genome sizes.

### Repetitive DNA sequences represent more than half of the *N. damascena* genome

For a more detailed analysis of genome and nuclear organization in giant-genome plants, we selected *N. damascena*, a plant with giant genome size (10.584 Gbp/1C; (Olszewska and Osiecka, 1983)). The karyotype of *N. damascena* has only 12 chromosomes (Natarajan and Ahnström, 1969; Klásterská and Natarajan, 1975; Orooji *et al*., 2022), suggesting that the large genome size is not determined by polyploidization but rather by the accumulation of repetitive DNA sequences. To estimate the content of repeats we performed next-generation DNA sequencing. Short, non-overlapping paired reads suitable for the RepeatExplorer2 assembly-free repeat analysis workflow were used to infer the repeat composition of the genome from 4,057,802 reads. About 58% of the reads belonged to 211 top clusters which were defined as those containing at least 405 reads (≥0.01% of analyzed reads) (Figure S4). Automated cluster annotation was manually verified, and for three clusters the annotations were corrected. The final verified cluster annotation was used for repeat quantification. 15 clusters containing 27,581 reads were annotated as organellar (plastid and mitochondrial) sequences. The remaining 4,030,221 reads represented nuclear DNA and corresponded to ∼0.06× coverage of the nuclear genome.

Circa 57% of the nuclear genome was composed of repetitive sequences, at which the repeatome was dominated by LTR-retrotransposons, representing ∼50% of the genome (Figure 2). The Tekay clade of chromoviral Ty3/Gypsy LTR-retrotransposons was the most abundant, comprising ∼30% of the genome. Other abundant LTR families included Ty3/Gypsy Athila (8.16%) and Ty3/Gypsy Retand (2.11%). Unclassified LTR elements and unclassified repeats occupied 8.54% and 5.54%, respectively, of the genome. DNA transposons (class II), including MuDR_Mutator and EnSpm_CACTA families, contributed only to 0.44% of the genome. Satellite DNA and rDNA corresponded to 0.47% and 0.32% of the genome coverage, respectively.

**Figure 2.**
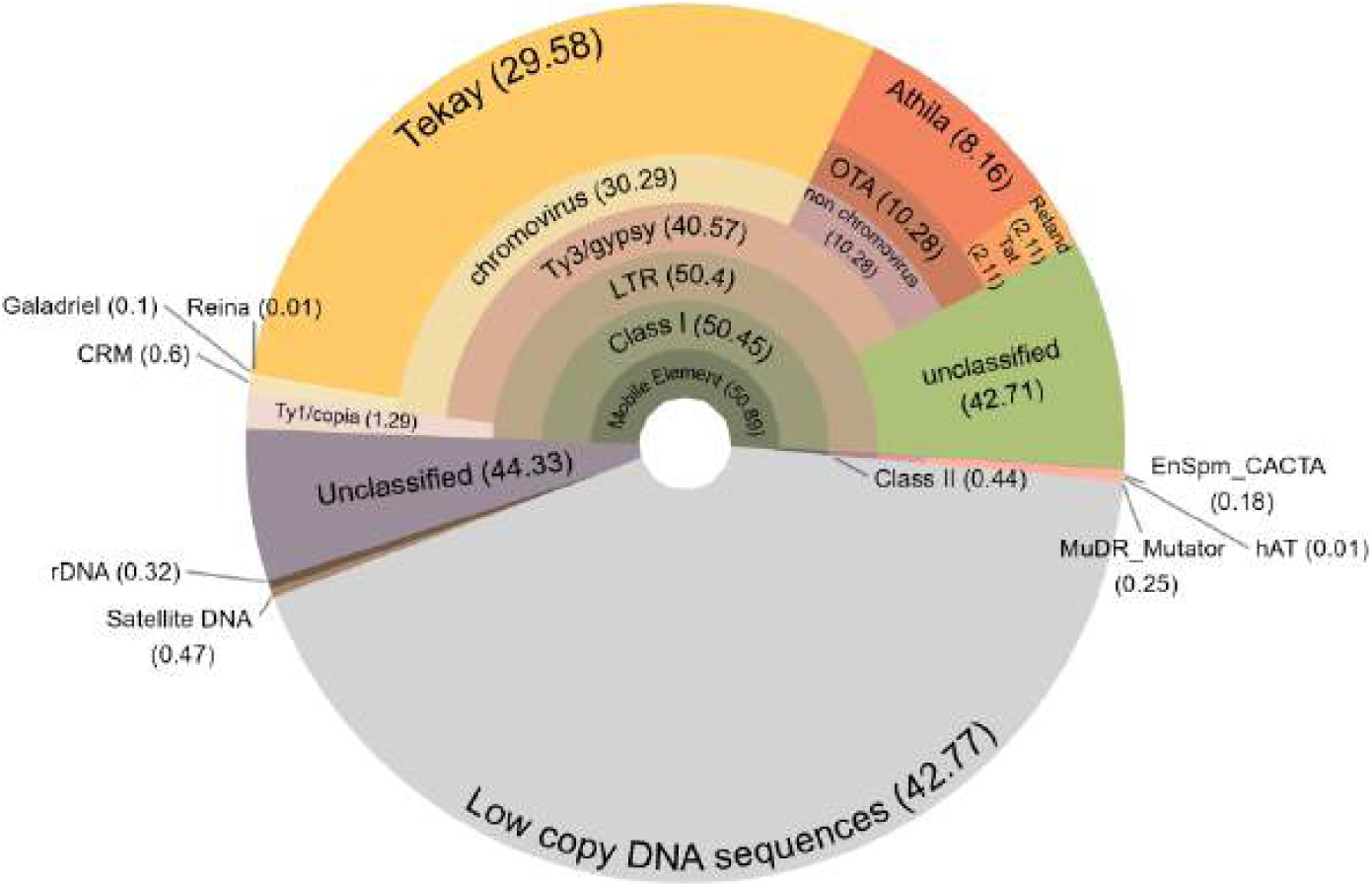
Types of high and moderate repeated DNA sequences in the *N. damascena* genome. The proportion of each repeat type or family is shown in the parentheses. Repeat proportions were calculated by annotating clusters representing at least 0.01% of the genome in RepeatExplorer2. For mobile elements, the members of each repeat family are indicated in the outer layers.

Thus, it seems plausible that the genome size of *N. damascena* is specifically related to the abundance of repeats in its composition.

### DNA replication kinetics in *N. damascena*

For the detection of replicating chromatin, we used 5-ethynyl-2′-deoxyuridine (EdU) which was incorporated into growing roots. In this study, we used several types of preparations that allowed us to analyze different aspects of DNA replication kinetics. First, we isolated nuclei from *N. damascena* roots and studied the localization of replicating chromatin using confocal microscopy. We identified three major staining patterns, differing by number and localization of EdU signals (Figure 3a). Type 1 nuclei contained many discrete EdU signals, which were preferentially distributed between DAPI-positive heterochromatin complexes. Type 2 nuclei exhibited significantly more intense labeling, with the label occupying a substantial portion of the nucleus volume and partially overlapping with heterochromatin. On the other hand, type 3 nuclei displayed only a few discrete signals that were associated with heterochromatin. It appears that these three types of nuclei may correspond to different S phase stages.

**Figure 3.**
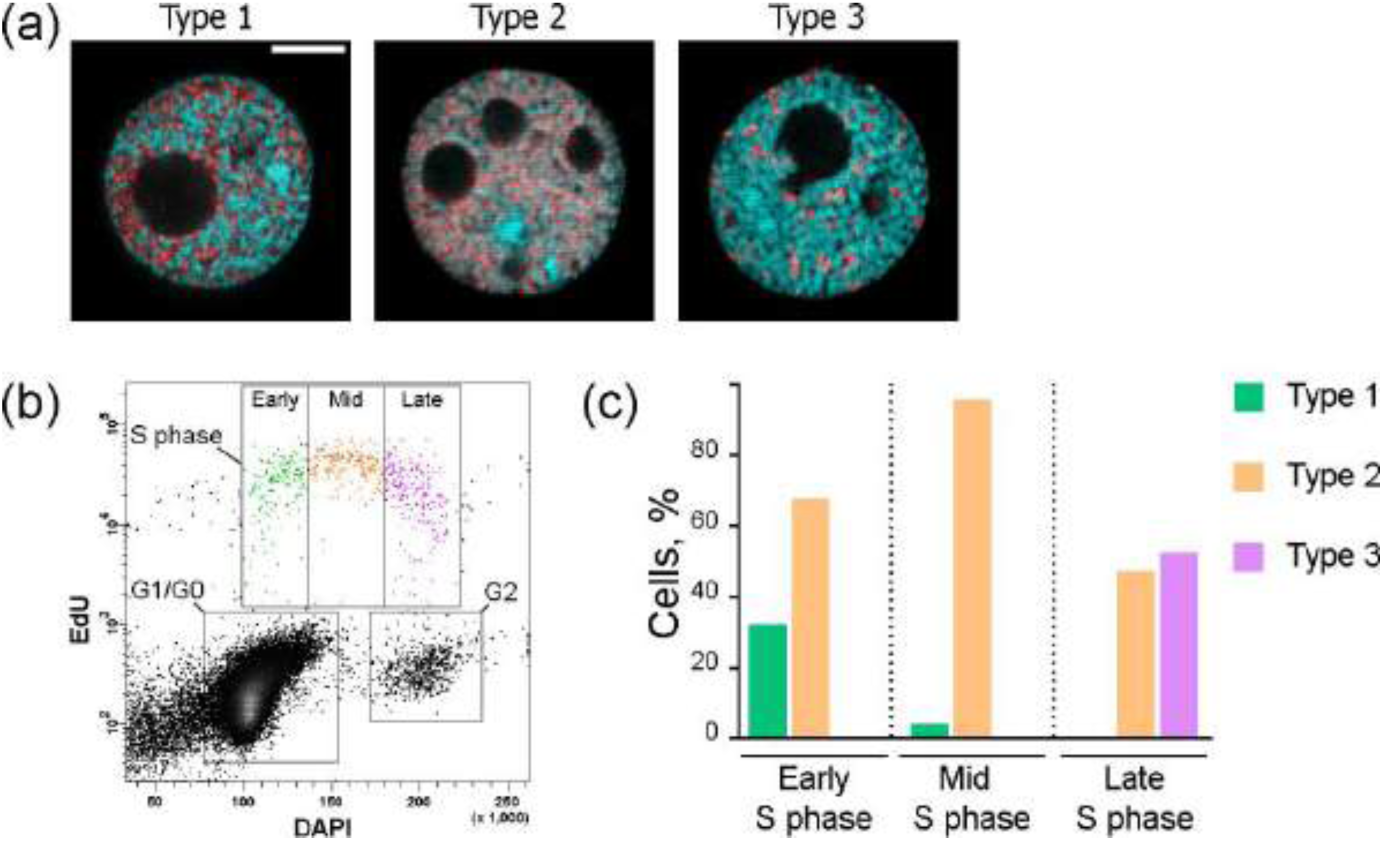
Chromatin replication patterns in meristematic root nuclei of *N. damascena*. (a) Three major patterns of EdU incorporation in isolated nuclei. Bar = 5 μm. (b) Flow cytometric analysis of the cell cycle. Bivariate analysis of EdU versus DAPI resulted in a typical pattern that allowed to differentiate between nuclei in G1/G0, S, and G2 stages. S phase cells were separated into three gates containing early, mid, and late S phase nuclei. (c) The S phase stages are characterized by different amounts of the three EdU incorporation patterns. Altogether 233 flow-sorted nuclei were analyzed by confocal microscopy (22 type 1, 157 type 2 and 54 type 3 nuclei).

Precise identification of replication stages was performed by sorting isolated root nuclei in which replication was detected by EdU. Bivariate flow cytometric analysis of EdU versus DAPI fluorescence enabled the distinction of nuclei in the G1 and G2 phases of the cell cycle, as well as those in the early, mid, and late S phases (Figure 3B). The intensity of EdU labeling was highest in mid S phase. Early, mid, and late S phase nuclei were separated and their morphology was analyzed (Figure 3c). Type 1 nuclei were more abundant in the fraction of early replication, type 2 nuclei in the fraction of mid replication, and type 3 nuclei in the fraction of late replication. With 67% of all S phase nuclei type 2 nuclei were the most numerous, and evident in all three fractions, indicating that the mid S phase was longer than both other stages (Figure 3c). Thus, it was possible to distinguish early, mid, and late S phase nuclei via EdU labeling.

### Replication of eu- and heterochromatin

When working with isolated nuclei, the impact of the effect of out-of-focus fluorescence on the images obtained was significant, even when using confocal microscopy (Figure 3a). To get rid of it, we prepared semithin (200-250 nm) sections of roots embedded in an acrylic medium (Figure 4a). EdU signals were concentrated in discrete foci within the euchromatin in early S phase nuclei. The nuclei in mid S phase were intensively labeled by EdU throughout the nucleoplasm, except for large blocks of condensed chromatin and nucleoli. The staining intensity in late S phase nuclei was substantially higher than in early S phase nuclei, making it easier to identify these patterns more reliably (Figure S5). Only a few distinct foci were visible within the nuclei during the late S phase.

**Figure 4.**
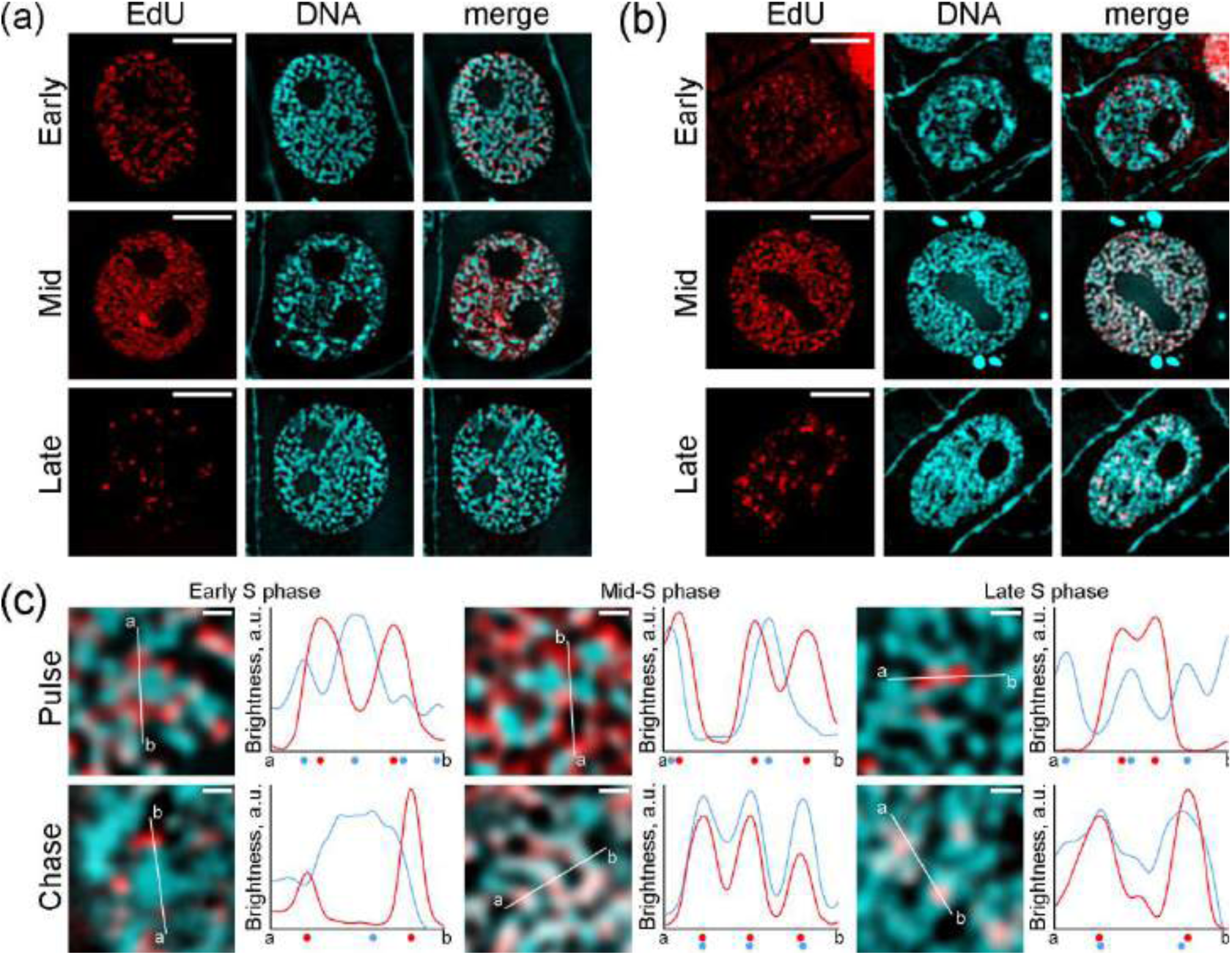
Replication kinetics in *N. damascena* root meristem cells. (a) Three labeling variants were classified under pulse conditions (labeling of chromatin during replication). For the acquisition of early S phase nuclei images, a 2-3 times higher exposure time was applied, required due to lower fluorescence intensity (see Figure S4). The chromonema organization was partially disassembled in mid S phase. Bar = 5 μm. (b) Three labeling types were classified under chase conditions (fixation of chromatin 2h after pulse labeling). The chromonemata in type 1 nuclei were decondensed, possibly because they were in mid S phase. Similarly, nuclei labeled with EdU in late S phase had a modified structure under chase conditions, possibly because some of them were in preprophase or even prophase. Bar = 5 μm. (c) Localization of replication foci (replicating chromatin) relative to DAPI-labeled chromonemata (heterochromatin) in all three nuclei types under pulse and chase conditions. Under pulse conditions the replicating chromatin is located always close to heterochromatin. Under chase conditions, the EdU labels were located similarly at the chromonemata surface in type 1 nuclei, or within the heterochromatin in type 2 and 3 nuclei. The dots under the density graphs indicate the positions of the maxima of heterochromatin (cyan) and EdU-containing chromatin (red). Bar = 0.5 μm (c).

The replication patterns observed in *N. damascena* differ significantly from those observed in mammals, in which, replication foci are distributed throughout the nucleoplasm, in the early S phase, discrete replication foci are observed in contact with nucleoli and the nuclear envelope, indicating facultative heterochromatin replication, and only a few discrete foci remain in late S phase, indicating constitutive heterochromatin replication (O’Keefe *et al*., 1992) (see also Figure S6). Therefore, it appears that the replication kinetics in *N. damascena* differs from those typically observed in animals. Obviously, mid S phase nuclei of *N. damascena* have no equivalent in animals, and vice versa.

In all three *N. damascena* replication patterns, the EdU signals colocalized with decondensed chromatin. As a result, it was difficult to determine whether euchromatin or heterochromatin was under replication. Therefore, the EdU labeling was additionally analyzed under chase conditions, i.e., the roots were fixed 2 h after pulse EdU incorporation, when replication was complete (Figure 4b). In early S phase nuclei, the distribution of EdU-labeled regions was not altered under chase conditions compared to pulse conditions. However, the mid and late S phase nuclei underwent a significant reorganization. The EdU was incorporated within the chromonemata during the mid S phase or within discrete heterochromatin blocks during the late S phase.

To provide further clarification on these visual observations, density profiles were established (Figure 4c). They indicate that after pulse labeling, the EdU signals are present within decondensed chromatin in all three nuclei types. However, after chase labeling, the signals were found within decondensed chromatin only in the early S phase, while they were located within condensed chromatin in the mid and late S phase nuclei. Thus, it appears that the heterochromatin underwent a decondensation during mid and late S phase replication, followed by a subsequent condensation when replication finished. Additionally, it was observed that the chromonemata were replicated during the mid S phase.

### Reorganization of the chromonema meshwork during replication

During the analysis of replication kinetics, it was observed that the distribution of chromonemata throughout the nucleoplasm was almost uniform in most nuclei. However, in the mid S phase nuclei, this uniform distribution was severely rearranged (Figure 3C, middle panels; Figure S7). In such nuclei, thick anastomosing strands of condensed chromatin (chromonemata) were still visible in some regions of the nucleoplasm, but in between the chromonemata also small chromatin complexes, such as globules and thin threads, appeared. Thus, the organization of condensed chromatin is heterogeneous during mid S phase, thus clearly distinguishing such nuclei from those in any other interphase stages. In these latter nuclei, the condensed chromatin forms a homogeneous meshwork of thick anastomosing chromonemata. As described above, the EdU signals were detected only in regions of decondensed chromatin, but not within chromonemata. The decondensation of chromonemata during replication significantly altered the global nuclear organization. Consequently, nuclei in mid of S phase could be identified by DAPI staining due to the absence of the regular meshwork of chromonemata.

The use of conventional light microscopy (LM) did not provide sufficient details to study the structure of the decondensation zones during the mid S phase, even when using semi-thin sections. To achieve more accurate visualization of chromonema reorganization during replication, we applied super-resolution 3D-SIM on semi-thin (∼200 nm) sections (Figure 5A). We performed both visual image analysis and density profile analysis to reveal the colocalization between condensed chromatin and replication domains. In early S phase nuclei, the replication foci were observed within decondensed chromatin, particularly in proximity to either chromonemata or sporadic thinner (∼100 nm) threads (Figure 5B, top panels). In mid S phase nuclei, 3D-SIM detected two variants of condensed chromatin: thick threads (i.e., interphase chromonemata) with a diameter of ∼300 nm, and small globules or chains of globules with a diameter of ∼100 nm. The replication foci were visible on the surface of interphase chromonemata, and close to small globules and chaines of condensed chromatin (Figure 5B, middle panels; Movie S1). Only in rare cases replication foci were not in contact with condensed chromatin. The nuclei of late S-phase contained two distinct types of replication loci: small foci and larger complexes, which seemed to be composed of multiple smaller foci (Figure 5B, bottom panels). Both of these types of replication complexes were in contact with chromonemata.

**Figure 5.**
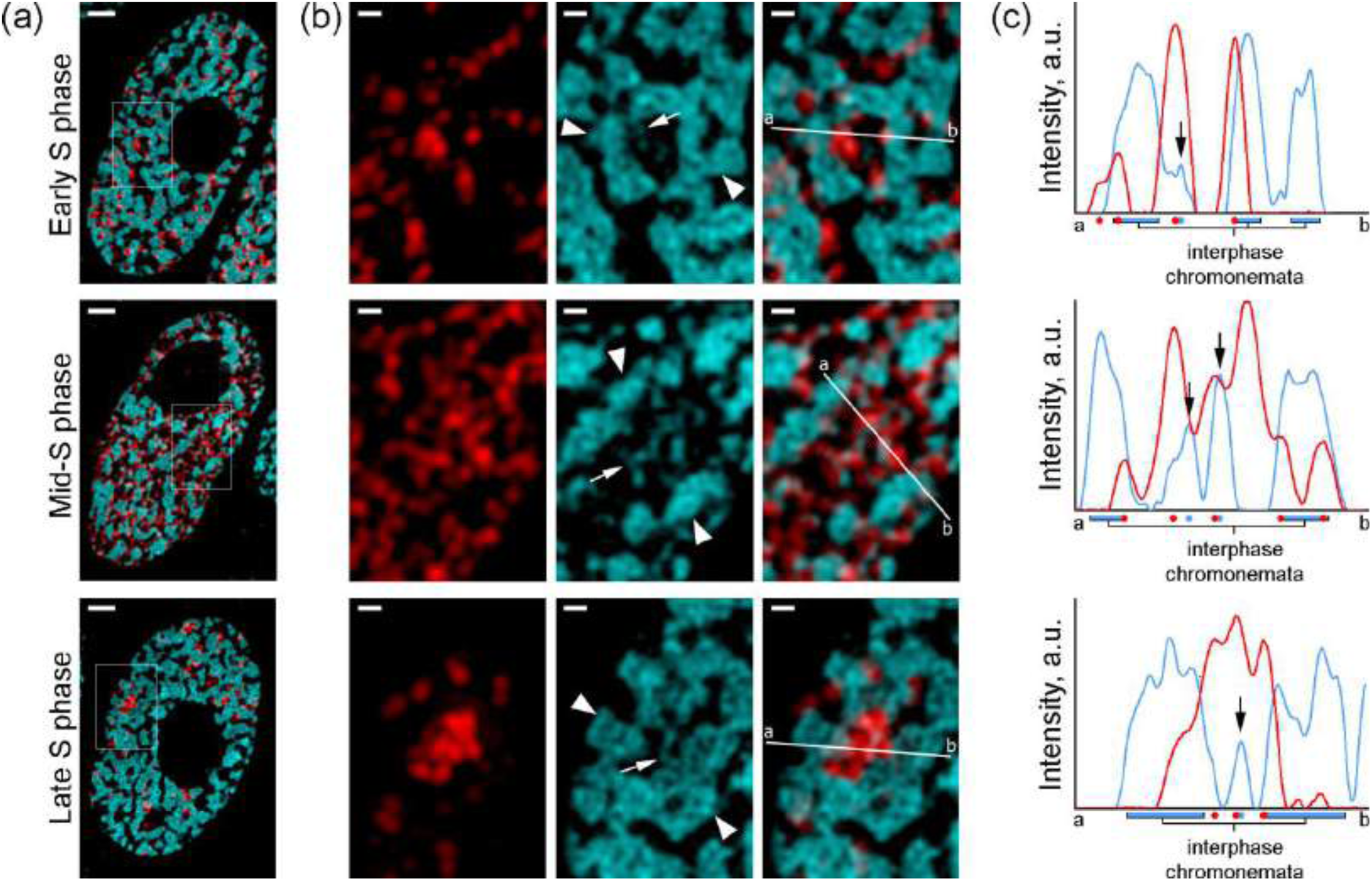
Ultrastructure of *N. damascena* nuclei during replication revealed via super-resolution 3D-SIM. (a) Early, mid and late S phase nuclei. (b) Enlarged regions indicated by rectangles in (a). Chromonemata are marked by arrowheads, thin threads of condensed chromatin by arrows. Chromonemata are indicated by arrowheads, thin threads of condensed chromatin by arrows. (c) Intensity plot profiles through the a-b lines in panel (b). In all cases, replicating chromatin (indicated by red dots under the plots) is located either at the boundary of thick threads of condensed chromatin (interphase chromonemata) or near thin threads of condensed chromatin (these threads are marked by blue dots under the plots and additionally indicated by black arrows in the plots).

During S-phase, DNA replication occurs mainly outside of condensed chromatin, but in contact with it. Replication causes the significant decondensation of chromonemata during mid S-phase, resulting in a global chromatin reorganization in interphase nuclei visible even in DAPI images. Nevertheless, despite the use of super-resolution microscopy, the details of replication zone organization could not be fully resolved. Therefore, also TEM was performed.

### Ultrastructural organization of replicating chromonemata

The CLEM approach, which combines LM and TEM, was used to study the ultrastructural organization of replicating chromatin (Figure S8A). A two-step non-rigid image registration approach, which was specifically developed for the alignment of LM and TEM images for correlative analysis, was used (Figure S8B). The images were registered using a contour-based approach presented by Sorokin et al. (2018), followed by a keypoint-based registration with mutual information as the similarity metric.

Due to the lower brightness of fluorescence images in ultrathin sections, which were used for CLEM analysis, compared to semithin sections, early S phase nuclei could not be analyzed. As a result, CLEM was used exclusively to analyze heterochromatin decondensation during replication and condensation after replication, specifically during mid and late S phase.

In the nucleoplasm of mid S phase nuclei, some regions had a morphological organization typical for *N. damascena* nuclei (extended thick chromonemata), but other regions contained numerous small (50-100 nm in diameter) discrete heterochromatin complexes of irregular shape that overlapped with EdU labeling under pulse conditions (Figure 6A). This suggests that these regions correspond to chromonema decondensation regions visible by LM (Figures 3C and 5B). At least partially, the EdU signals overlapped with heterochromatin (Figure 6A; Figure S9). In contrast, under chase conditions, heterochromatin was preferentially organized into thick threads (chromonemata), and EdU labeling preferentially overlapped with heterochromatin (Figure 6B).

**Figure 6.**
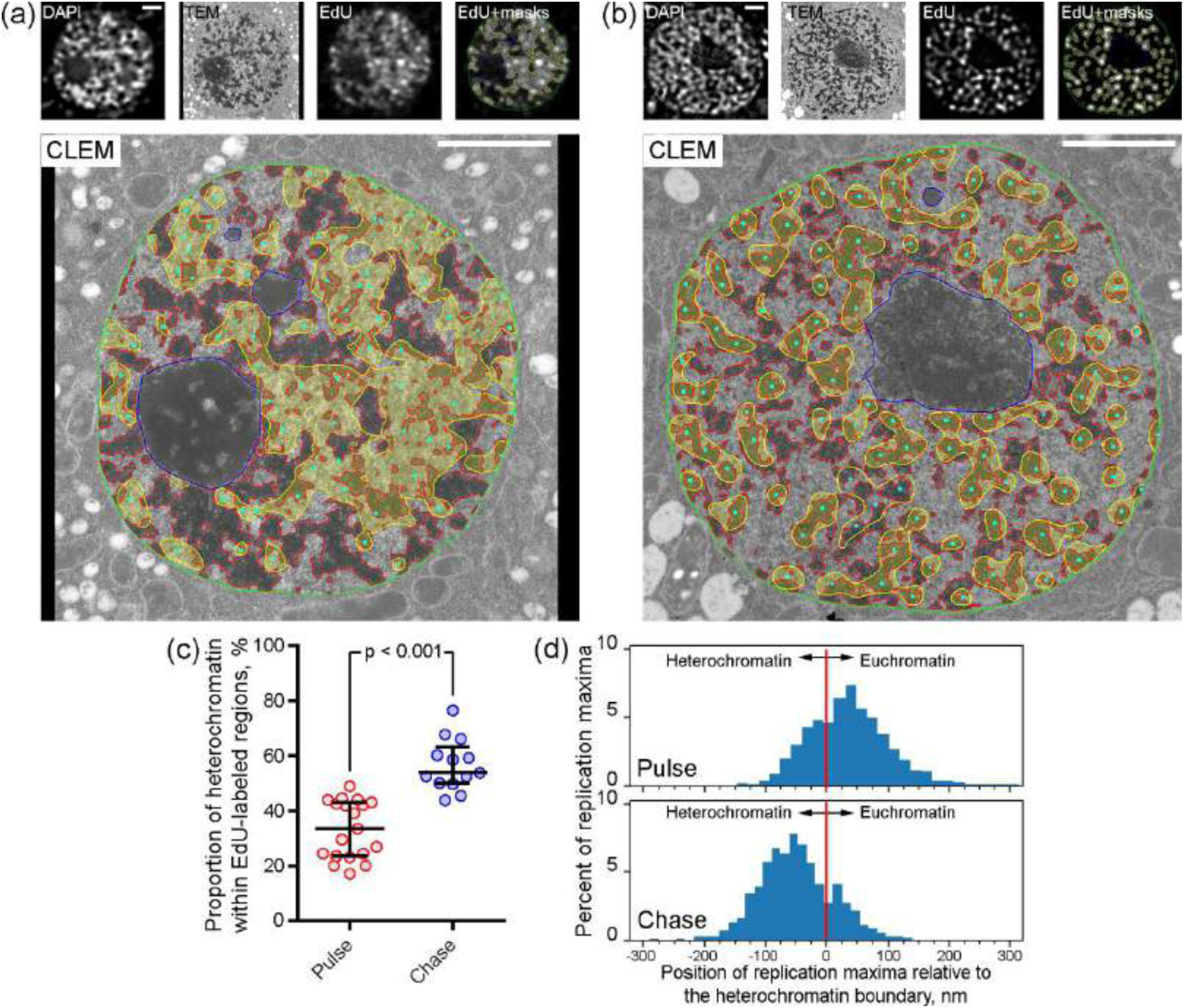
Correlative light and electron microscopy (CLEM) of replicating chromatin in *N. damascena* mid S phase nuclei. (a) CLEM of a nucleus labeled by EdU under pulse conditions (labeling of replicating chromatin). The images of the top panel were used for the segmentation of the nuclear structures shown in the bottom panel (blue contours - nucleoli; red contours – heterochromatin; yellow contours and filling - replication complexes; cyan dots - centroids of replication complexes). The replication complexes overlap preferentially with the decondensed chromatin compartments containing small heterochromatin complexes. Bars = 2 μm. (b) CLEM of a nucleus labeled by EdU under chase conditions (labeling of chromatin, whose replication was finished before fixation). The images of the top panel were used for the segmentation of the nuclear structures (color markings as in a). The replication complexes overlap preferentially with heterochromatin, which forms extended chromonemata. Bars = 2 μm. (c) Estimation of the overlap between replication complexes and heterochromatin. The proportion of the area of regions labeled with EdU, occupied by heterochromatin, is larger under chase than pulse conditions. The comparison was performed using the Mann–Whitney U test. (d) The localization of replication complex centroids relative to heterochromatin borders. Under pulse conditions, the majority of centroids localized in the euchromatin compartments, but under chase conditions, the majority of centroids located inside heterochromatin.

Using CLEM, we measured the colocalization of replicating chromatin in LM images with condensed chromatin in TEM images. We segmented replicating chromatin in EdU images and condensed chromatin in TEM images using adaptive thresholding techniques and measured the relative overlap of the resulting segmentation masks. We found that the overlap was increased under chase conditions compared to pulse conditions (Figure 6C). However, the fluorescence masks had relatively low resolution. Therefore, we additionally calculated the centroids of replication spots as the local maxima of the denoised image intensity, assuming that these spots are closest to the replicating chromatin (cyan dots in Figures 6A and B). Then, we estimated the centroid positions relative to the boundary between heterochromatin and the interchromatin compartment. Replication centroids were preferentially located in the euchromatin compartments under pulse conditions, and in heterochromatin under chase conditions (Figure 6D). The rare labeling of heterochromatin complexes in the pulse experiments could be the result of the rapid recondensation of chromatin after replication (to increase the labeling intensity, we incubated the roots in EdU for a relatively long time of 30 min, and some chromatin complexes could become recondensed already during the labeling period).

A similar relocation of EdU signals was observed in late S phase nuclei. The EdU pulse labels appeared to be concentrated within decondensed chromatin. In contrast to mid S phase nuclei, the label was concentrated in a smaller number of foci, and the chromonemata organization was not altered (Figure S10A). As in middle S phase nuclei, chase EdU signals overlapped with condensed chromatin (Figure S10B).

## DISCUSSION

The profound variation in genome size between species has been described in several plants and animals, which has occurred preferentially through the expansion of different non-coding DNA sequences (Gregory, 2005). Plants with giant genomes are of particular interest because an excess of DNA can lead to changes in the organization of interphase nuclei and mitotic chromosomes (Kuznetsova *et al*., 2017). For instance, growing genome sizes during the evolution of eukaryotes required dense chromatin compaction realized via the helical organization of sister chromatids in metaphase chromosomes larger than ∼12 Mb (Câmara, et al., 2023). In early TEM investigations on plant chromatin ultrastructure, networks of thick heterochromatin filaments were proven in interphase nuclei of *Tradescantia paludosa*, and these filaments were termed ‘chromonemata’ (Sparvoli *et al*., 1965). To our knowledge, such a heterochromatin organization has never been observed in animal nuclei.

While mammalian genome sizes vary only slightly (Ryan Gregory, 2011; Redi and Capanna, 2012; Kapusta *et al*., 2017), there is tremendous variability in genome size and complexity among different plant species (Michael, 2014; Pellicer and Leitch, 2020). Among Angiosperms, genome sizes vary remarkably by > 2,200-fold, ranging from 67.2 Mbp/1C in *Genlisea aurea* (Fleischmann *et al*., 2014) to 149.185 Gbp/1C in *Paris japonica* (Pellicer *et al*., 2010). Variation in genome size in green plants is often due to whole-genome duplications, which are widespread in plants (Van de Peer *et al*., 2017; Li *et al*., 2021), and on the transposable element gains and losses (Lee and Kim, 2014). Here, we show that three previously identified variants of plant nuclei: chromocentric, chromomeric, and chromonemal (Nagl *et al*., 1983), were found in the nuclei of plants with small (<1.5 Gbp/1C), large (∼1.5- 5 Gbp), and giant (>5 Gbp) genomes, respectively. Thus, genome expansion alters the morphology of plant interphase nuclei by increasing the amount of condensed chromatin and altering its localization.

Our main goal was to investigate the less studied chromonemal variant of heterochromatin organization typical of giant genome plants. For a detailed investigation, we selected *N. damascena*, a common weed species in the Euro-Mediterranean region and a popular garden plant known as love-in-a-mist. *N. damascena* has a giant genome size of 10.584 Gbp/1C (Olszewska and Osiecka, 1983), which is about 24 times larger than the genome of the model plant *Arabidopsis thaliana* (0.435 Gbp/1C; Schmuths et al. 2004). Using NGS sequencing, we showed that the *N. damascena* genome contains numerous repeats (preferentially LTR retrotransposons) representing >50% of the global genome. Heterochromatin in *N. damascena* nuclei formed typical chromonemata that filled the entire volume of the interphase nuclei except the nucleoli.

Heterochromatin can undergo cycles of decompaction-compaction during replication, which has been studied in detail in cultured mammalian cells (Chagin *et al*., 2019), but DNA replication kinetics in plants appear to be different from those described in animal cells. The early S phase in plants is characterized by weakly dispersed replication signals, the speckled signals concentrated in specific areas were typical for the late S phase, whereas strong signals dispersed throughout the nucleus except for the nucleolar area were observed in the mid S phase nuclei (Jasencakova *et al*., 2001; Bass *et al*., 2015; Němečková *et al*., 2020). Such patterns were found in *Hordeum vulgare* L. (Jasencakova *et al*., 2001; Bass *et al*., 2015; Němečková *et al*., 2020), *Triticum aestivum* L., *Avena sativa* L., *Secale cereale* L., *Oryza sativa* L., *Brachypodium distachyon* L., (Jasencakova *et al*., 2001; Bass *et al*., 2015; Němečková *et al*., 2020), *Zea mays* L. (Jasencakova *et al*., 2001; Bass *et al*., 2015; Němečková *et al*., 2020), and our results in *N. damascena* were consistent with these data. It seems that such replication kinetics are typical for plants, at least for plants with large and giant genomes. However, to describe the fine organization of chromonemata during and after replication, it was necessary to apply more sophisticated microscopic methods to resolve at the ultrastructural level.

Previously, we have proposed the use of semi-thin sections of roots embedded in acrylic resin to identify replicating chromatin (Kuznetsova *et al*., 2017; Sheval, 2018). This method allows the examination of extremely thin sections (200-250 nm) with a 3-4 times higher axial resolution than reached by confocal microscopy. Using this approach, we were able to identify precisely the localization of replication sites relative to condensed chromatin. At all stages of S phase, signals were localized outside of condensed chromatin. Heterochromatin replicated in mid and late S phases, but it decondensed, and recondensed afterward.

The abundant amount of chromatin within chromonemata may also explain the high labeling intensity during mid S phase nuclei, which was clearly visible by both flow cytometry and LM. The consequence of this abundance was also a global reorganization of the nuclei: a substantial part of all chromonemata was decompacted, and thus mid S phase nuclei were even detectable by DAPI staining. In animal cells, only local and subtle heterochromatin reorganization in the S phase can be observed (Chagin *et al*., 2019). In contrast, gross chromatin reorganization during the S phase has been described in several papers (Lafontaine and Lord, 1974; de la Torre et al., 1975; Hao et al., 1994). However, all these studies were performed using autoradiography, a method that does not allow accurate identification of S-phase stages and monitoring of chromatin reorganization processes. Our data suggest that only the replication of dense chromonema chromatin leads to such a robust nuclear reorganization.

In addition, to characterize the organization of the replicating chromonemata (mid S nuclei), we used super-resolution 3D-SIM and CLEM microscopy, which allows us to detect the ultrastructural organization of chromatin. 3D-SIM showed that the replicating chromatin dispersed by decondensation is located close to the chromonemata. CLEM confirmed these observations: during replication, chromonemata were partially decondensed, but some small complexes of condensed chromatin were visible in the replication zones. The high resolution of TEM allowed us to precisely localize EdU signals within these zones, and we found that replication occurred outside these discrete remnants of chromonemata. On the contrary, after replication (chase conditions), the labels were localized almost entirely within the chromonemata (mid S nuclei) or blocks of condensed chromatin (late S nuclei).

In summary, we conclude that the replication of chromatin forming chromonemata in giant-genome plant nuclei is accompanied by a decompaction-compaction process similar to that described for animals. However, the abundance of such chromatin leads to a robust reorganization of the nuclear morphology not previously described in animals or plants with small genomes.

## MATERIALS AND METHODS

### Experimental objects

Seeds from 32 different species (Table 1) were cultivated for several days in a humidified chamber at 23°C protected from light. Most of the seeds were purchased from the Gavrish Company (Russia), except for *Hordeum vulgare* seeds, which were a generous gift from Prof. L.I. Khrustaleva. For *Vicia* species, we re-analyzed images previously obtained in our laboratory.

### Transmission electron microscopy (TEM)

Root tips were fixed in 2.5% glutaraldehyde in 100 mM cacodylate buffer, post-fixed in 1% OsO_4_, dehydrated in ethanol and propylene oxide, and embedded in SPI-Pon 812 (SPI Supplies, Cat. No. 02660-AB). Ultrathin sections (∼90 nm) were stained with uranyl acetate and lead citrate and examined with a JEM-1400 electron microscope (Jeol).

### DNA extraction and sequencing

Genomic DNA was extracted from the roots of *N. damascena* using the CTAB method (Doyle and Doyle, 1987). Library preparation followed the NEBNext Ultra II DNA Library Prep Kit for Illumina protocol (New England Biolabs). Size selection of adapter-ligated DNA was performed as described in the protocol, and the average length of the DNA library fragments was ∼400 bp. Genome sequencing was performed using the Illumina MiSeq system to obtain ∼20 mln paired-end (2×301 bp) reads. The genome size (in bp) of *N. damascena* was calculated using the formula: *genome size* = *DNA content (pg)* × (0.978 × 10^9^). The DNA content of *N. damascena* was determined to be 10.5 pg/1C (https://cvalues.science.kew.org/), and the genome size was estimated to be 10.269 Gbp/1C. The coverage of the sequenced genome was calculated according to the following formula: *coverage* = (*number of reads* × *read length*) / *1C genome size*.

### Repeat identification and annotation

Repetitive elements in the genome were analyzed using the graph-based clustering algorithm implemented in the RepeatExplorer2 pipeline (Novák *et al*., 2020). Raw DNA reads were uploaded to the Galaxy server with installed RepeatExplorer tools available at https://repeatexplorer-elixir.cerit-sc.cz. The quality of the sequencing reads was checked using FastQC installed on the server. The reads were trimmed to the length of 151 nucleotides and filtered by the quality of 95% of bases equal to or above the quality cutoff value of 10. The random sampling of 5 mln read pairs was conducted with RepeatExplorer utilities on the server. Paired-end read sequences were converted to FASTA format and interlaced into a single file. Only read pairs with no overlap were selected for the graph-based clustering analysis performed with the “RepeatExplorer2 clustering” tool. The automatic annotations of the discovered clusters of repetitive elements were manually corrected.

### Replication labeling

*N. damascena* L. seeds were purchased from the Gavrish Company (Cat. No. 002364). The seeds were grown in a Petri dish covered with filter paper at 25°С in the dark. Circa 10 mm long roots were used for the study. For pulse labeling, the germinated seeds were incubated in 50 μM EdU; (Invitrogen, Cat. No. A10044) for 15 min (CLEM) or 30 min (LM). In the case of chase labeling, EdU was then substituted with 200 μM thymidine for 30 min. Thymidine was removed subsequently and the germinated seeds were cultivated additionally for 1.5 hours in distilled water before fixation.

### Light microscopy

After EdU incorporation, 1.0 mm long root tips were excised and fixed with 2% formaldehyde in 0.5×PBS for 2 h. After fixation, they were embedded in LR White resin (Sigma, Cat. No. L9774). Semi-thin (∼200 nm) sections were cut using either an Ultratome LKB-III (LKB) or UltraCut E (Reichert-Jung). EdU was detected using a Click-iT EdU Cell Proliferation Kit for Imaging (Invitrogen, Cat. No. C10338). DNA was stained with DAPI (Invitrogen, Cat. No. D1306) and mounted in Mowiol 4-88 (Aldrich, Cat. No. 81381) containing the anti-bleaching agent DABCO (Sigma, Cat. No. D-2522). Sections were imaged using an Axiovert 200M microscope equipped with a 100×/1.4 Plan-Apochromat objective (Carl Zeiss GmbH) and a CCD camera ORCAII-ERG2 (Hamamatsu). Each LM experiment was performed twice, with a minimum of one hundred nuclei visually analyzed for each point. The non-S phase cells did not incorporate EdU and were used as a negative control.

To analyze the ultrastructure of chromatin and replication signals beyond the classical lateral Abbe-Rayleigh limit of ∼250 nm, spatial structured illumination microscopy (3D-SIM) was performed to achieve a lateral resolution of ∼140 nm (super-resolution, attained with a 561 nm laser). We used an Elyra PS.1 microscope system equipped with a 63×/1.4 Plan-Apochromat objective and the ZENBlack software (Carl Zeiss GmbH). Image stacks were captured separately for each fluorochrome using 405 nm (DAPI), and 561 nm (Alexa Fluor 555) laser lines for excitation and appropriate emission filters (Weisshart *et al*., 2016; Kubalová *et al*., 2021). Zoom-in sections are presented as single slices to detect the subnuclear chromatin structures at the super-resolution level.

### Fluorescence-activated sorting of isolated nuclei

Suspensions of intact nuclei were prepared, according to Guillotin et al. (2023) with minor modifications. Briefly, ∼0.5-1.5 cm long seedlings were fixed with 2% formaldehyde in 15 mM Tris buffer at 4°C and washed twice with Tris buffer at 4°C. Meristematic parts of roots (∼2 mm in length) were excised from ∼200 seedlings. Root meristems were homogenized in 2 ml pre-cooled lysis buffer (0.3 M sucrose, 15 mM Tris-HCl at pH 8, 60 mM KCl, 15 mM NaCl, 2 mM EDTA, 0.5 mM spermine, 0.5 mM spermidine, 15 mM MES, 0.1% Triton, 5 mM DTT, 0.4% bovine serum albumin, protease inhibitor cocktail 1:100 (Sigma, Cat. No. P8849), using a Dounce homogenizer. The resulting suspension was filtered through a 50 µm polyester filter (Filcons, Cat. No. 12050-47S) and centrifuged at 13000 g for 1 minute. The pellet was resuspended in 0.5 mL of Click-iT reaction cocktail prepared according to the manufacturer’s instructions, and the nuclei were incubated in the dark at 24°C for 60 minutes. The reaction cocktail was then washed with Tris buffer and the nuclei were stained with DAPI (1 µg/ml). Finally, the suspension of nuclei was analyzed using a FASCAria II SORP sorter (BD Biosciences) equipped with 405 and 561 nm lasers. Nuclei representing different phases of the cell cycle were sorted on a 24×60 polylysine-coated coverslip and mounted in Mowiol 4-88.

### Correlative light and electron microscopy (CLEM)

For CLEM, we adopted a previously published protocol (Arifulin, 2015). After EdU labeling, root tips were fixed in 2.5% glutaraldehyde in 100 mM cacodylate buffer overnight, followed by a brief permeabilization in 0.1% Triton X-100 for 5 min. For EdU detection, the Click-iT EdU Cell Proliferation Kit for Imaging, AlexaFluor 555 dye a Click-iT EdU Cell Proliferation Kit for Imaging was used, but the incubation in the reaction mixture was extended to 16 h. Subsequently, the samples were embedded in SPI-Pon 812 resin as described above. Ultrathin (∼100 nm) sections were cut on an Ultracut E ultramicrotome (Reichert-Jung) and mounted on single-slot grids with formvar mounting film. The sections were stained with 2 µg/ml DAPI in 1×PBS and then carefully mounted on a microscope slide under a coverslip in a drop of distillate water. The sections were photographed using a Nikon C2 microscope with a 60×/1.4 Plan-Apochromat objective. Then, the coverslips were removed, and the grids were stained with uranyl acetate and lead citrate and examined with a JEM-1400 electron microscope (Jeol).

Since some deformations and distortions may occur during this double imaging process, we developed special software to accurately superimpose LM and TEM images. First, we manually segmented the nucleus in both TEM and LM images. For LM, we used the DAPI channel because the nuclear structure appears more prominent. Then, we upscaled the LM images to the same resolution as the TEM images, from 1px = 63.4 nm to 1px = 4.4 nm. Since the LM images are quite smooth, such an upscaling did not introduce significant artifacts into the images.

Image registration was performed in two steps: coarse and fine. The coarse registration used the elasticity-based cell image registration method described elsewhere (Sorokin *et al*., 2018; Sorokin *et al*., 2020). The method uses the nuclear mask as a reference and transforms the LM images into the TEM images so that the nuclear masks are perfectly aligned. The transformation is constrained by elasticity laws and corresponds to the nature of the actual nucleus motion. Fine-tuning was performed using an intensity-based approach. The condensed chromatin areas in DAPI were segmented and used as key points for optimization-based image registration. We chose the thin-plate spline motion model with these key points as control points and mutual information as the cost function because the images are from different modalities. For both registration stages, we applied the same transformation to both channels of the LM images.

In round nuclei, registration resulted in almost perfect image overlap, but in elongated nuclei, we achieved only partial overlap. Therefore, we visually assessed the accuracy of the alignment of condensed chromatin images in the TEM and DAPI images and removed the regions with inaccurate alignment from the analysis. 19 nuclei were analyzed for pulse labeling, and 13 nuclei were analyzed for chase labeling.

## ACKNOWLEDGMENTS

We are grateful to A.V. Lazarev for technical support, A.A. Penin and L.I. Khrustaleva for advice and discussion. Computational resources for RepeatExplorer analysis were provided by the ELIXIR-CZ project (LM2023055), part of the international ELIXIR infrastructure. This work was supported by the Russian Foundation for Basic Research (20-54-12016) and by the Deutsche Forschungsgemeinschaft (Schu 762/12-1).

## AUTHOR CONTRIBUTIONS

EVS, TDK and VS, conceived the study and designed the experiments; EAA, MAK, DMP, DOO, and VS performed most of the experiments; DVS, and NAA performed image analysis; AAV performed the data analysis; EAA, DVS, AAV, VS, and EVS wrote the manuscript.

## CONFLICT OF INTEREST

The authors declare that they have no competing interests.

## DATA AVAILABILITY STATEMENT

Raw DNA sequencing data were deposited in Sequence Read Archive (SRA) with accession number PRJNA939363. Code to reproduce the CLEM analysis is provided at https://gitlab.com/dsorokin.msk/clem-image-analysis and https://gitlab.com/dsorokin.msk/mmb-ipnb-tracking. The other data that support the findings of this study are available from the corresponding authors upon reasonable request.

## Supplementary Figures

**Figure S1.**
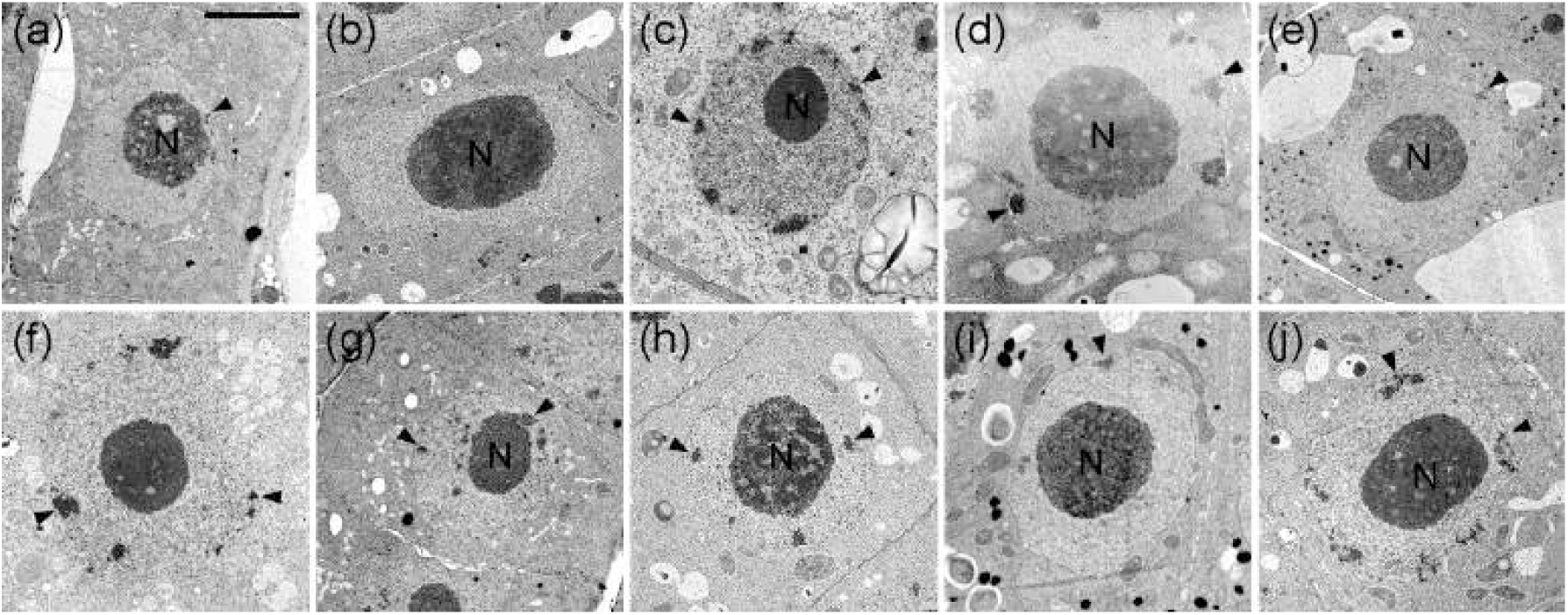
Representative nuclei of species with low heterochromatin content. Heterochromatin forms discrete complexes, chromocenters (arrowheads). Nucleoli are indicated by (N). (a) *Daucus carota* L.; (b) *Brassica rapa* L.; (c) *Cucurbita pepo* L.; (d) *Linum usitatissimum* L.; (e) *Brassica oleracea* L.; (f) *Phaseolus vulgaris* L.; (g) *Catharanthus roseus* (L.) G.Don; (h) *Lupinus polyphyllus* Lindl.; (i) *Cucumis sativus* L.; (j) *Solanum lycopersicum* L. Bar = 5 μm.

**Figure S2.**
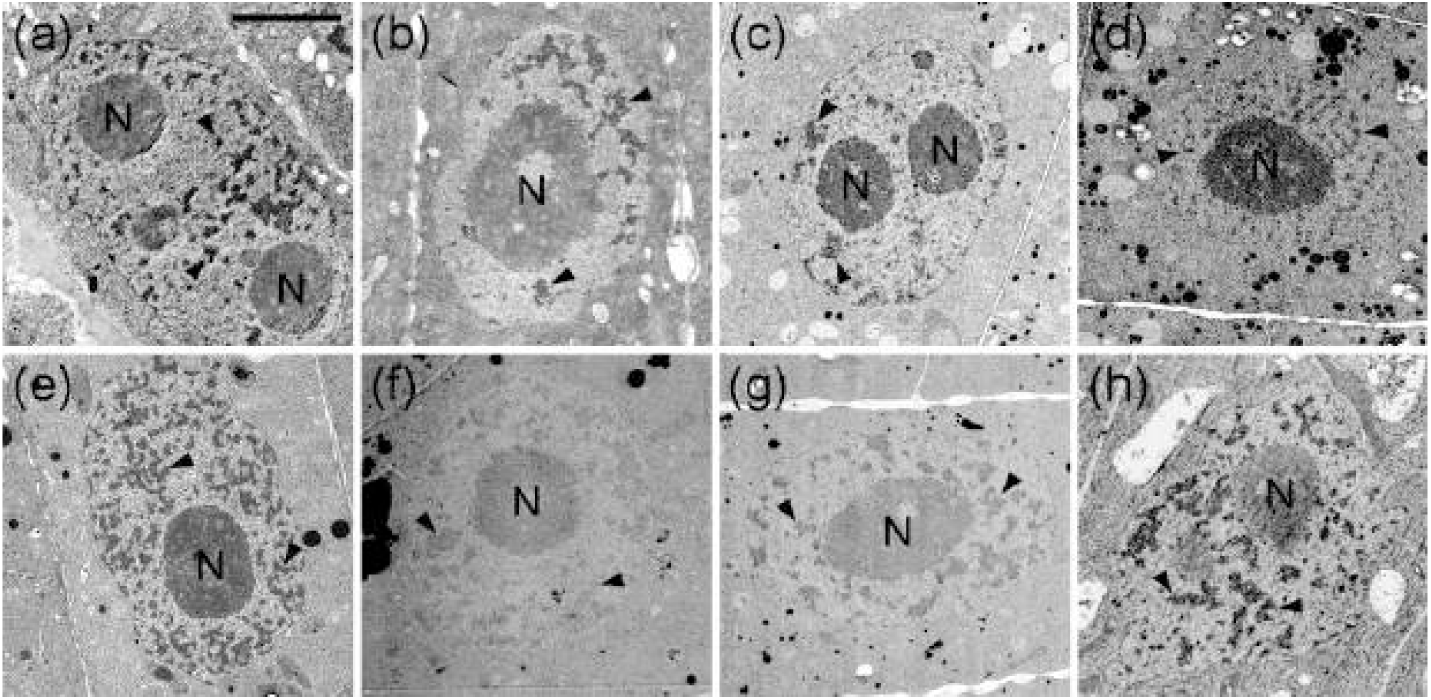
Representative nuclei of species with a moderate amount of heterochromatin. Heterochromatin forms discrete complexes, chromomeres (arrowheads), which fill the entire nucleoplasm. Nucleoli are indicated by (N). (a) *Daucus carota* L.; (b) *Brassica rapa* L.; (c) *Cucurbita pepo* L.; (d) *Linum usitatissimum* L.; (e) *Brassica oleracea* L.; (f) *Phaseolus vulgaris* L.; (g) *Catharanthus roseus* (L.) G.Don; (h) *Lupinus polyphyllus* Lindl.; (i) *Cucumis sativus* L.; (j) *Solanum lycopersicum* L. Bar = 5 μm.

**Figure S3.**
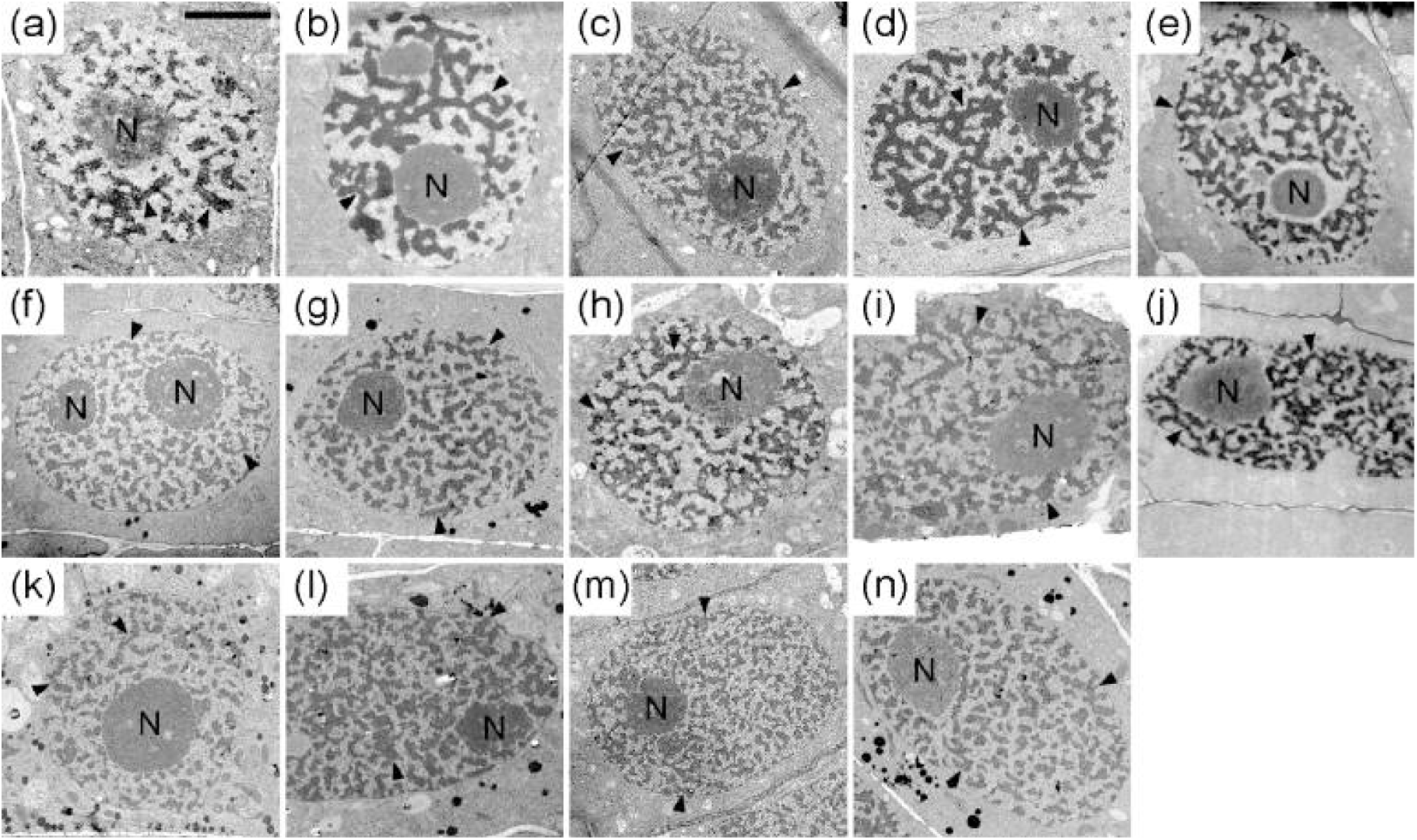
Representative nuclei of species with a high heterochromatin amount. Heterochromatin forms thick threads, chromonemata (arrowheads), which fill the entire nucleoplasm. Nucleoli are indicated by (N). (a) *Vicia sepium* L.; (b) *Hordeum vulgare* L.; (c) *Lathyrus odoratus* L.; (d) *Lathyrus latifolius* L.; (e) *Nigella sativa* L.; (f) *Nigella damascena* L.; (g) *Allium fistulosum* L.; (h) *Vicia faba* L.; (i) *Allium altaicum* Pall.; (j) *Allium sativum* L.; (k) *Allium schoenoprasum* L.; (l) *Allium cepa* L.; (m) *Allium porrum* L.; (n) *Allium ramosum* L. Bar = 5 μm.

**Figure S4.**
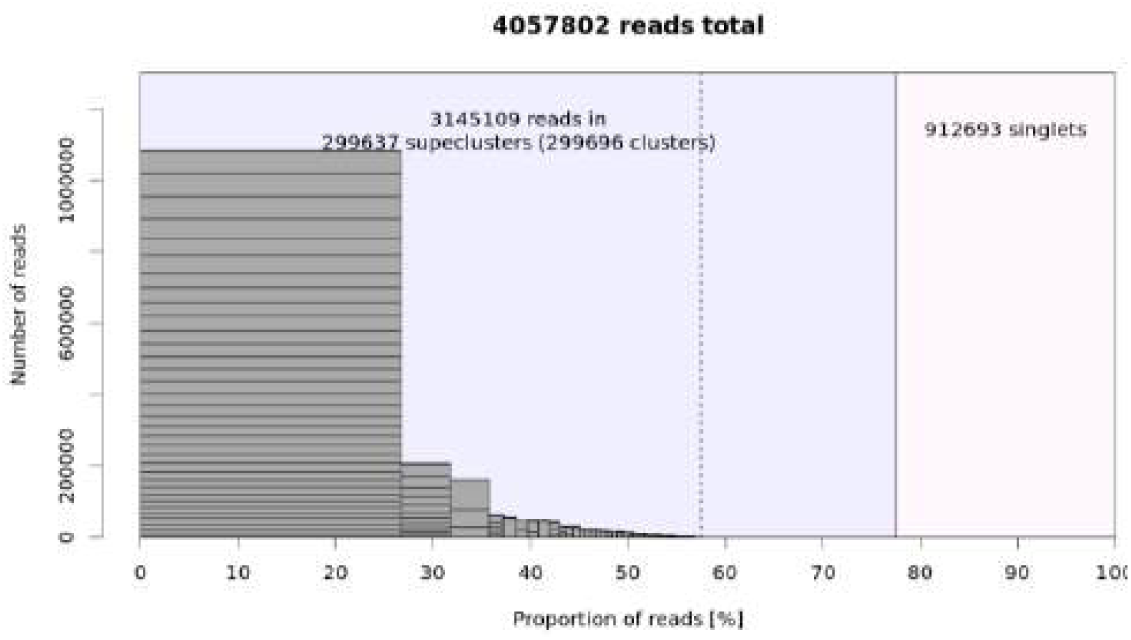
Graphical summary of the RepeatExplorer2 clustering results. Bars represent superclusters, and rectangles inside bars represent individual clusters. Top clusters are shown on the left of the dotted line.

**Figure S5.**
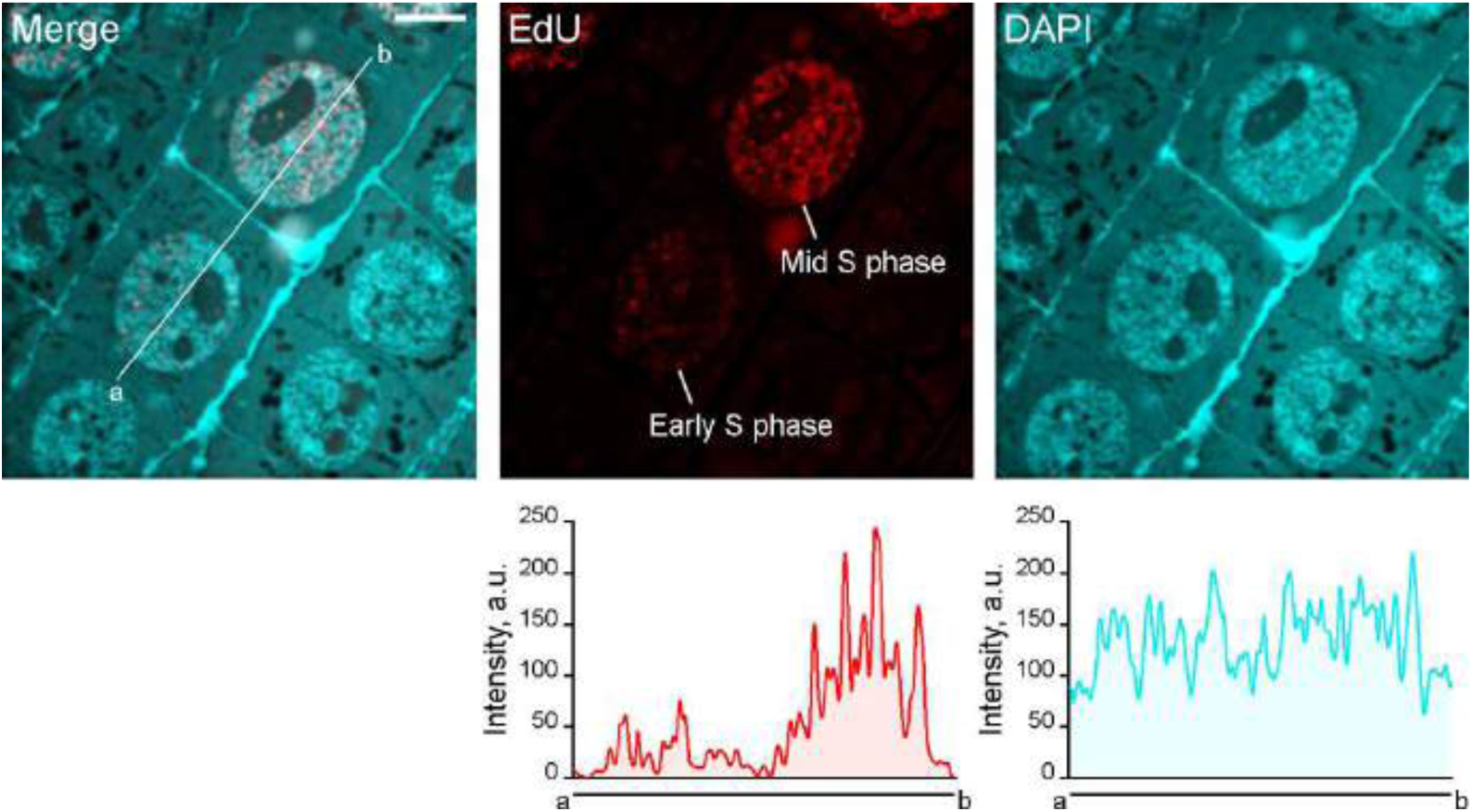
EdU labeling of *N. damascena* S phase nuclei. EdU labeling is significantly brighter in the mid S phase than in the early S phase, as also visible in the brightness diagrams. The images were not processed to preserve the true brightness relationships between different cells. Bar = 5 μm.

**Figure S6.**
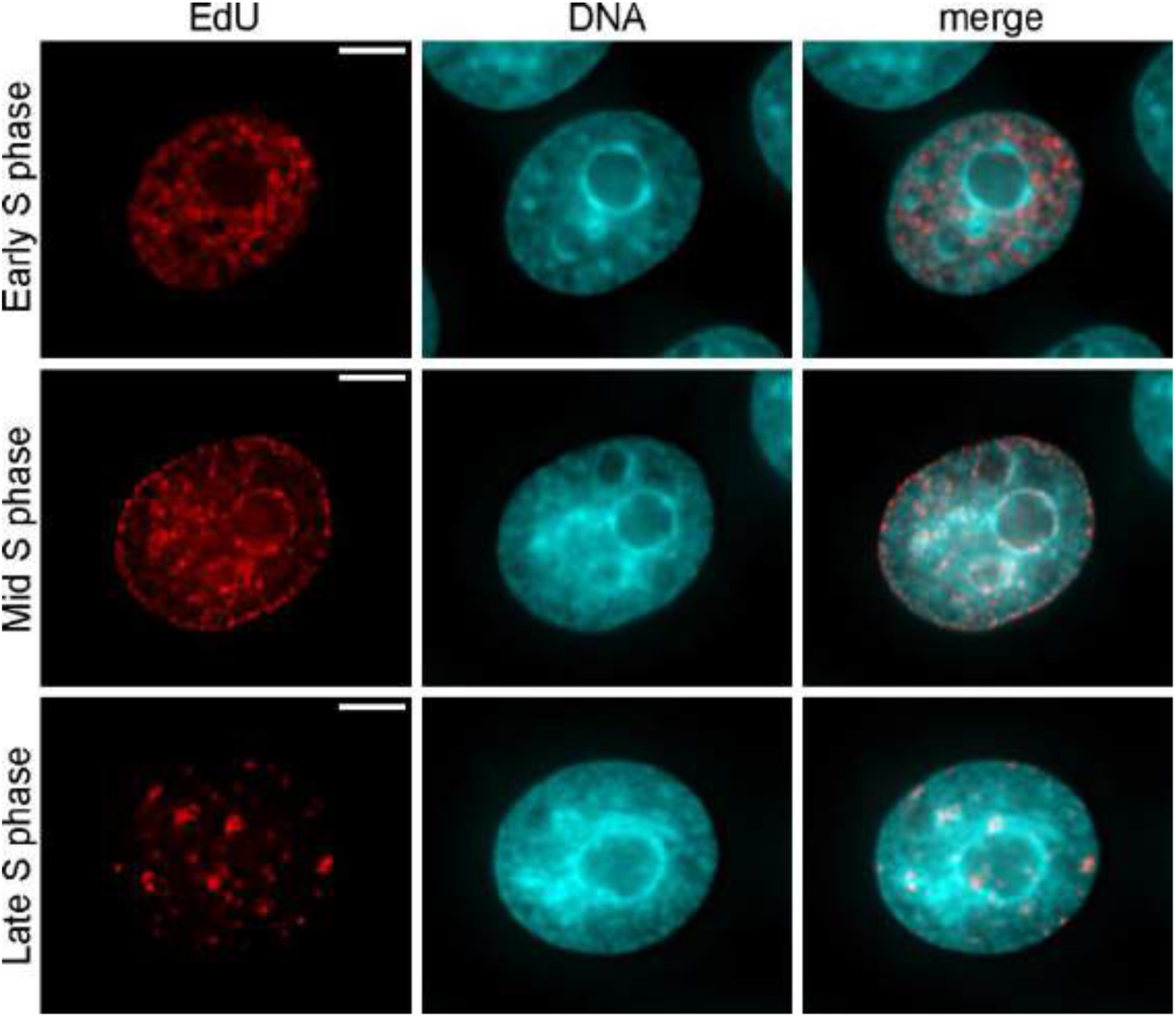
Replication kinetics in human carcinoma HeLa cells. During the early S phase, the euchromatin replication foci are distributed evenly throughout the nucleoplasm and are absent in heterochromatin. In mid S phase, discrete replication foci are visible preferentially in contact with nucleoli and the nuclear envelope (facultative heterochromatin replication). In the late S phase, only a few discrete foci remain (constitutive heterochromatin replication). Such a kinetics is typical of many cultured cells in different mammals. Bars = 5 μm.

**Figure S7.**
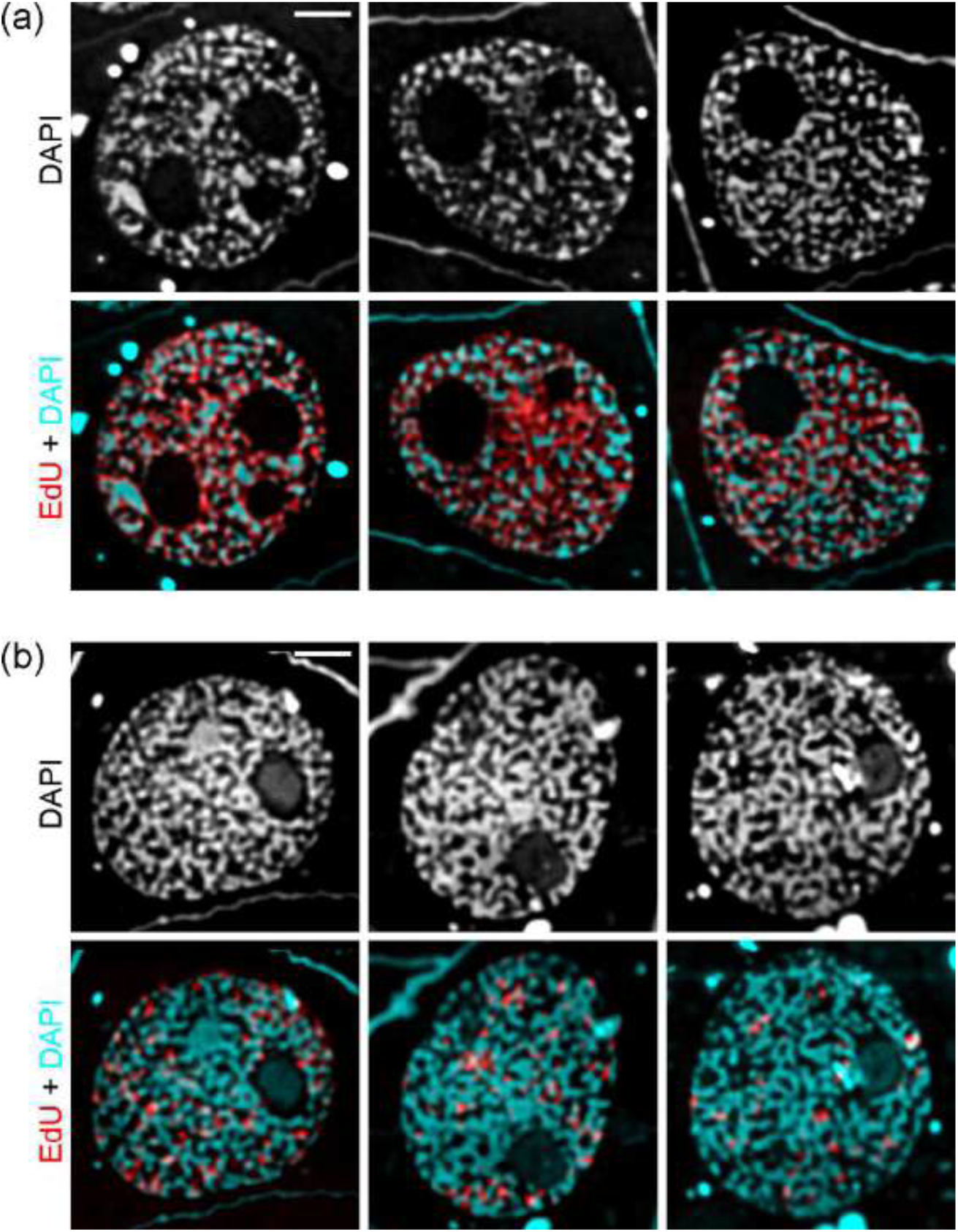
Chromonema meshwork reorganization occurs in *N. damascena* mid S phase nuclei. Compared to mid S phase (a) the majority of late S phase nuclei (b) show highly condensed chromatin forming thick anastomosing threads (chromonemata). This meshwork fills the entire interior of the interphase nuclei. In most mid S phase nuclei, the condensed chromatin forms predominantly discrete globular or elongated complexes of obviously different sizes. This allows for clearly distinguishing mid S phase from other stages, even when only stained by DAPI. Bars = 2 μm.

**Figure S8.**
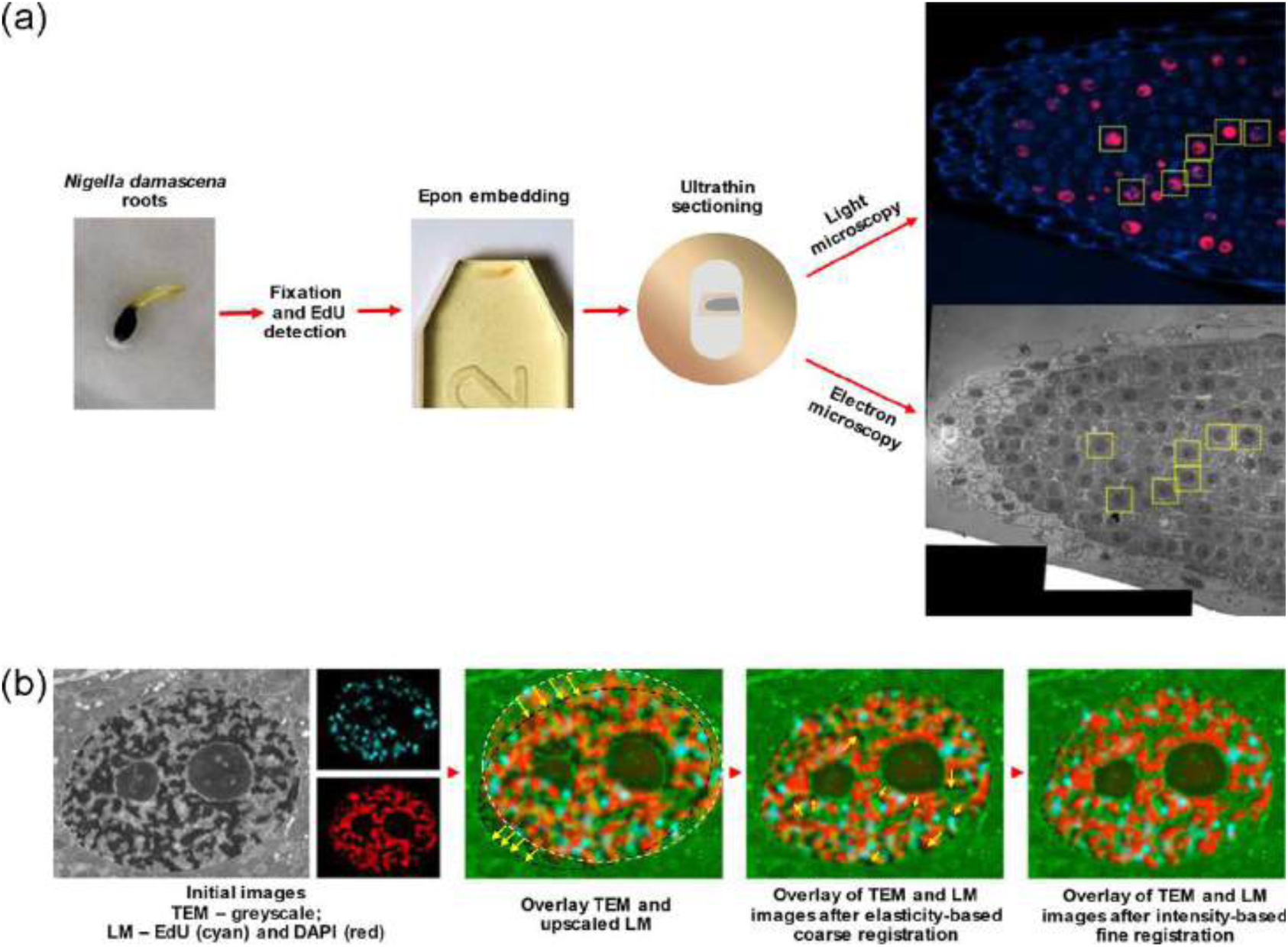
Major steps of correlative light and electron microscopy (CLEM) in *N. damascena*. (a) The seeds were grown in a Petri dish covered with filter paper at 25°С in the dark. The germinated seeds were incubated in EdU for 15 min. After EdU labeling, root tips were fixed in 2.5% glutaraldehyde, followed by EdU detection. Subsequently, the samples were embedded in SPI-Pon 812 resin. Ultrathin (∼100 nm) sections were mounted on single-slot grids with formvar mounting film. The sections were stained with DAPI and then carefully mounted on a microscope slide under a coverslip in a drop of distilled water. The sections were photographed using a fluorescence microscope. Then, the coverslips were removed, and the grids were stained with uranyl acetate and lead citrate. Finally, the grids were examined with a transmission electron microscope. (b) The initial fluorescence and electron microscopy images and LM images in original resolution with their segmentation masks. The upscaled LM images (in cyan and red) overlayed with the TEM image (in gray). The top panels show the DAPI channel, bottom panels show the EdU channel. The overlay of the LM and TEM images after the coarse elasticity-based stage of the registration. The overlay of the LM and TEM images after the fine intensity-based stage of the registration.

**Figure S9.**
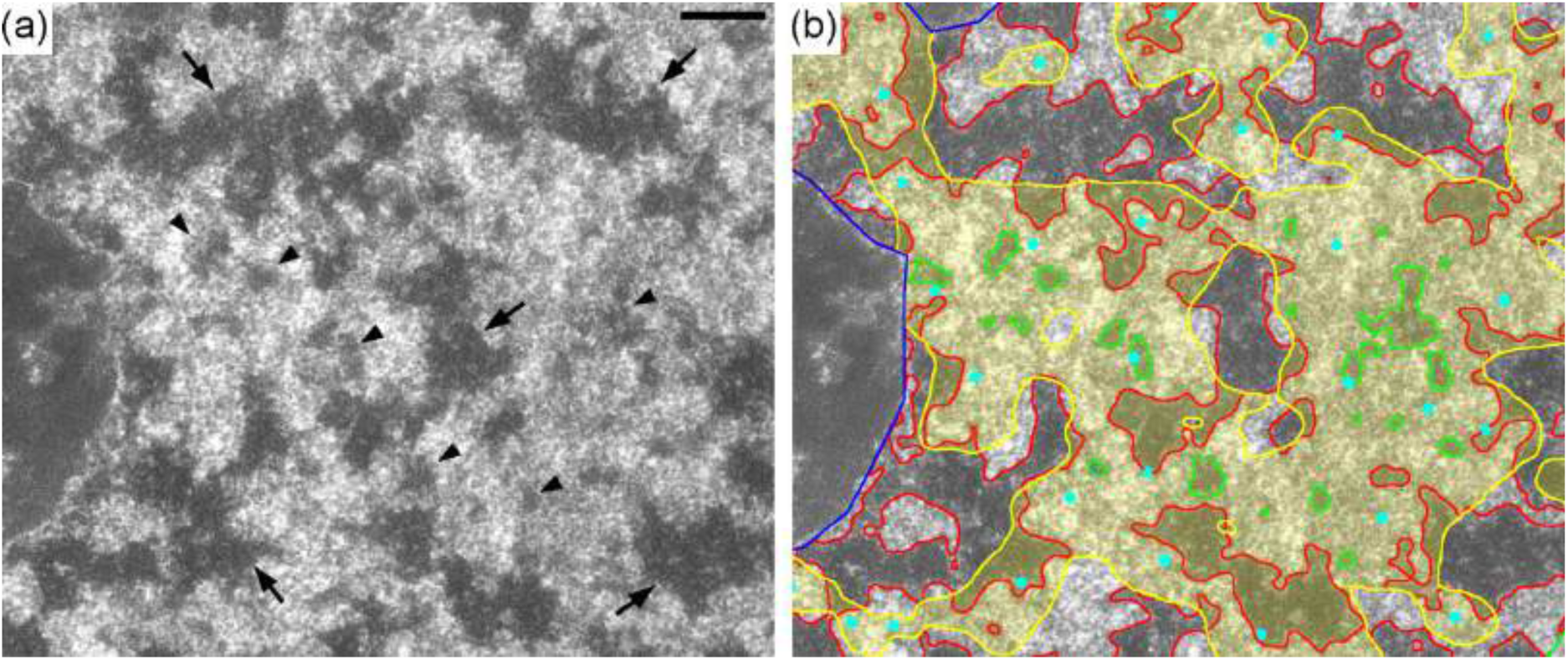
CLEM of replicating chromatin in *N. damascena* mid S phase nuclei. (a) Fragment of the nucleus shown in Figure 5A. The chromonemata are partially decondensed. Chromonemata are indicated by arrows; discrete chromatin complexes between chromonemata (the result of chromonema decondensation) are indicated by arrowheads. (b) The result of segmentation (blue contours - nucleoli, red contours - heterochromatin (chromonemata), yellow contours and filling - EdU signals, cyan dots - centroids of EdU signals). The replication complexes preferentially overlap the interchromatin compartment where small heterochromatin complexes (green contours) are visible. Bar = 0.5 μm.

**Figure S10.**
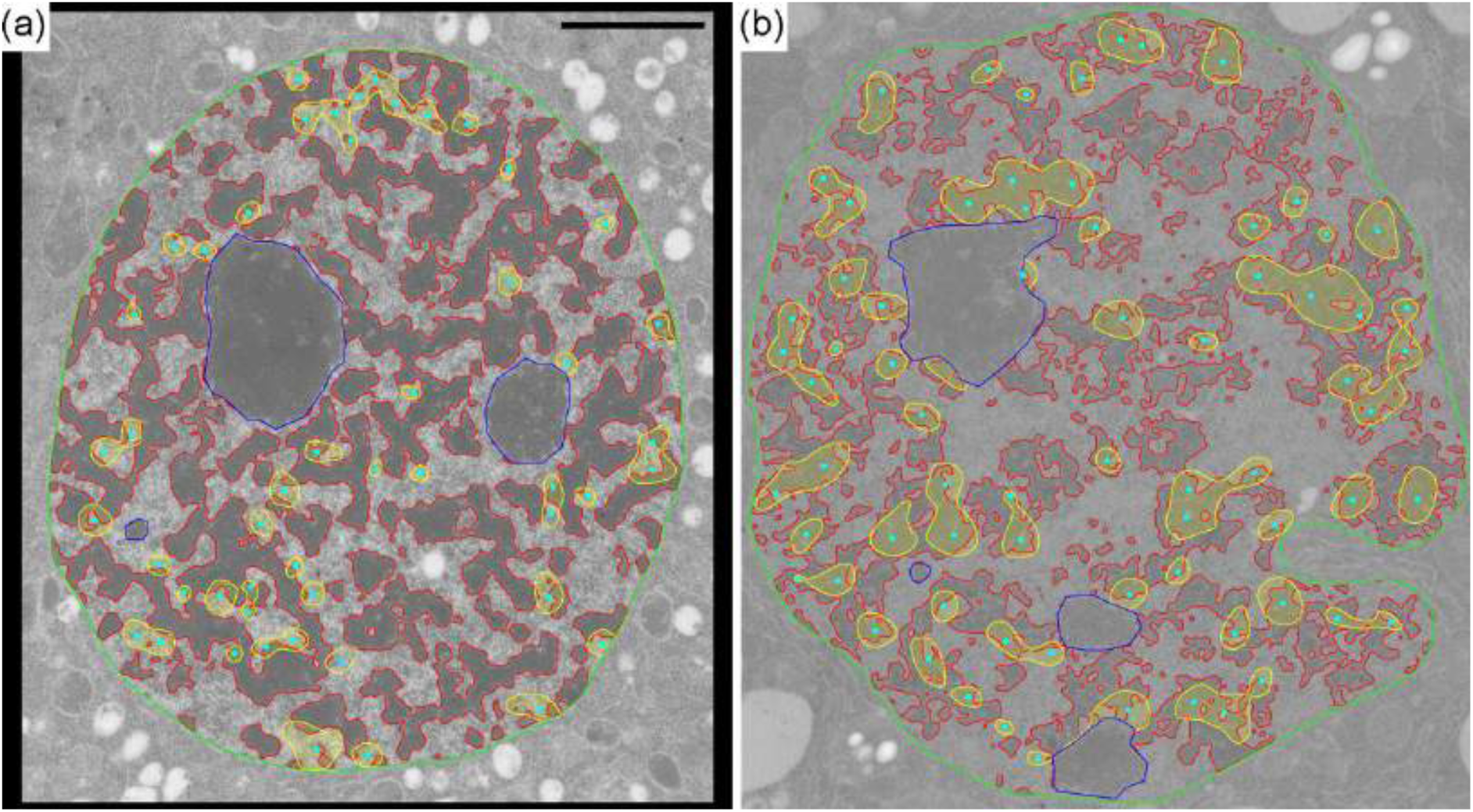
CLEM of replicating chromatin in *N. damascena* late S phase nuclei. (a) Nucleus labeled with EdU under pulse conditions (fixation of just replicating chromatin). The replication complexes overlap preferentially with decondensed chromatin compartments. (b) Nucleus labeled with EdU under chase conditions (fixation of chromatin, whose replication was finished before fixation). Replication complexes overlap with condensed chromatin. It should be emphasized that condensed chromatin is substantially reorganized, probably due to the onset of prophase chromosome condensation. For segmentation color description, see Figure S9. Bar = 2 μm.

## Notes

### Competing Interest Statement

The authors have declared no competing interest.

### Summary of Updates

Text and figures were revised. Authors list was changed.

## References

Arifulin, E.A. (2015) Ultrastructural organization of replicating chromatin in prematurely condensed chromosomes. Biopolymers and Cell, 31, 249–254.

Arumuganathan, K. & Earle, E.D. (1991) Nuclear DNA content of some important plant species. Plant Molecular Biology Reporter, 9, 208–218.

Bai, C., Alverson, W.S., Follansbee, A. & Waller, D.M. (2012) New reports of nuclear DNA content for 407 vascular plant taxa from the United States. Annals of Botany, 110, 1623–1629.

Baranetzky, J. (1880) Die Kerntheilung in den Pollenmutterzellen einiger Tradescantien. Botanische Zeitschrift, 38 281–296.

Baranyi, M. & Greilhuber, J. (1999) Genome size in *Allium*: in quest of reproducible data. Annals of Botany, 83, 687–695.

Barow, M. & Meister, A. (2003) Endopolyploidy in seed plants is differently correlated to systematics, organ, life strategy and genome size. *Plant*, Cell & Environment, 26, 571– 584.

Bass, H.W., Hoffman, G.G., Lee, T.-J., Wear, E.E., Joseph, S.R., Allen, G.C., Hanley-Bowdoin, L. & Thompson, W.F. (2015) Defining multiple, distinct, and shared spatiotemporal patterns of DNA replication and endoreduplication from 3D image analysis of developing maize (*Zea mays* L.) root tip nuclei. Plant Molecular Biology, 89, 339–351.

Bennett, M.D. & Smith, J.B. (1976) Nuclear DNA amounts in angiosperms. Philosophical Transactions of the Royal Society B Biological Sciences, 274, 227–274.

Câmara, A.S., Kubalová, I. & Schubert, V. (2023) Helical chromonema coiling is conserved in eukaryotes. The Plant Journal, 118, 1284–1300

Ceccarelli, M., Morosi, L. & Cionini, P.G. (1998) Chromocenter association in plant cell nuclei: determinants, functional significance, and evolutionary implications. Genome, 41, 96–103.

Chagin, V.O., Reinhart, B., Becker, A., Mortusewicz, O., Jost, K.L., Rapp, A., Leonhardt, H. & Cardoso, M.C. (2019) Processive DNA synthesis is associated with localized decompaction of constitutive heterochromatin at the sites of DNA replication and repair. Nucleus, 10, 231–253.

Chen, M.-S., Niu, L., Zhao, M.-L., et al. (2020) De novo genome assembly and Hi-C analysis reveal an association between chromatin architecture alterations and sex differentiation in the woody plant *Jatropha curcas*. Gigascience, 9, giaa009.

Dong, P., Tu, X., Chu, P.-Y., Lü, P., Zhu, N., Grierson, D., Du, B., Li, P. & Zhong, S. (2017) 3D chromatin architecture of large plant genomes determined by local A/B compartments. Molecular Plant, 10, 1497–1509.

Doyle, J.J. & Doyle, J.L. (1987) A rapid DNA isolation procedure for small quantities of fresh leaf tissue. Phytochemical Bulletin, 19, 11–15.

Falk, M., Feodorova, Y., Naumova, N., et al. (2019) Heterochromatin drives compartmentalization of inverted and conventional nuclei. Nature, 570, 395–399.

Feodorova, Y., Falk, M., Mirny, L.A. & Solovei, I. (2020) Viewing nuclear architecture through the eyes of nocturnal mammals. Trends in Cell Biology, 30, 276–289.

Fleischmann, A., Michael, T.P., Rivadavia, F., Sousa, A., Wang, W., Temsch, E.M., Greilhuber, J., Müller, K.F. & Heubl, G. (2014) Evolution of genome size and chromosome number in the carnivorous plant genus *Genlisea* (Lentibulariaceae), with a new estimate of the minimum genome size in angiosperms. Annals of Botany, 114, 1651–1663.

Fransz, P., de Jong, J.H., Lysak, M., Castiglione, M.R. & Schubert, I. (2002) Interphase chromosomes in *Arabidopsis* are organized as well defined chromocenters from which euchromatin loops emanate. Proceedings of the National Academy of Sciences of the United States of America, 99, 14584–14589.

Fransz, P. & de Jong, H. (2011) From nucleosome to chromosome: a dynamic organization of genetic information. The Plant Journal, 66, 4–17.

Gregory, T.R. (2005) The C-value enigma in plants and animals: a review of parallels and an appeal for partnership. Annals of Botany, 95, 133–146.

Grime, J.P. & Mowforth, M.A. (1982) Variation in genome size – an ecological interpretation. Nature, 299, 151–153.

Guillotin, B., Rahni, R., Passalacqua, M., et al. (2023) A pan-grass transcriptome reveals patterns of cellular divergence in crops. Nature, 617, 785–791.

Hao, S., Jiao, M., Zhao, J., Xing, M. & Huang, B. (1994) Reorganization and condensation of chromatin in mitotic prophase nuclei of *Allium cepa*. Chromosoma, 103, 432–440.

Hendrix, B. & Stewart, J.M. (2005) Estimation of the nuclear DNA content of gossypium species. Annals of Botany, 95, 789–797.

Jakob, S.S., Meister, A. & Blattner, F.R. (2004) The considerable genome size variation of *Hordeum* species (Poaceae) is linked to phylogeny, life form, ecology, and speciation rates. Molecular Biology and Evolution, 21, 860–869.

Jasencakova, Z., Meister, A. & Schubert, I. (2001) Chromatin organization and its relation to replication and histone acetylation during the cell cycle in barley. Chromosoma, 110, 83–92.

Johnston, J.S., Pepper, A.E., Hall, A.E., Chen, Z.J., Hodnett, G., Drabek, J., Lopez, R. & Price, H.J. (2005) Evolution of genome size in Brassicaceae. Annals of Botany, 95, 229–235.

Kapusta, A., Suh, A. & Feschotte, C. (2017) Dynamics of genome size evolution in birds and mammals. Proceedings of the National Academy of Sciences of the United States of America, 114, E1460–E1469.

Klásterská, I. & Natarajan, A.T. (1975) Distribution of heterochromatin in the chromosomes of *Nigella damascena* and *Vicia faba*. Hereditas, 79, 154–156.

Kron, P. & Husband, B.C. (2012) Using flow cytometry to estimate pollen DNA content: improved methodology and applications. Annals of Botany, 110, 1067–1078.

Kubalová, I., Câmara, A.S., Cápal, P., et al. (2023) Helical coiling of metaphase chromatids. Nucleic Acids Research, 51, 2641–2654.

Kubalová, I., Němečková, A., Weisshart, K., Hřibová, E. & Schubert, V. (2021) Comparing super-resolution microscopy techniques to analyze chromosomes. International Journal of Molecular Sciences, 22, 1903.

Kubešová, M., Moravcova, L., Suda, J., Jarošík, V. & Pyšek, P. (2010) Naturalized plants have smaller genomes than their non-invading relatives: a flow cytometric analysis of the Czech alien flora. Preslia, 82, 81–96.

Kuznetsova, M.A., Chaban, I.A. & Sheval, E.V. (2017) Visualization of chromosome condensation in plants with large chromosomes. BMC Plant Biology, 17, 153.

Kuznetsova, M.A. & Sheval, E.V. (2016) Chromatin fibers: from classical descriptions to modern interpretation. Cell Biology International, 40, 1140–1151.

Lafontaine, J.G. & Lord, A. (1974) An ultrastructural and radioautographic study of the evolution of the interphase nucleus in plant meristematic cells (*Allium porrum*). Journal of Cell Science, 14, 263–287.

Lee, S.-I. & Kim, N.-S. (2014) Transposable elements and genome size variations in plants. Genomics & Informatics, 12, 87–97.

Leitch, I.J., Hanson, L., Lim, K.Y., Kovarik, A., Chase, M.W., Clarkson, J.J. & Leitch, A.R. (2008) The ups and downs of genome size evolution in polyploid species of *Nicotiana* (Solanaceae). Annals of Bot., 101, 805–814.

Li, B., Lin, D., Zhai, X., et al. (2022) Conformational changes in three-dimensional chromatin structure in *Paulownia fortunei* after Phytoplasma infection. Phytopathology, 112, 373–386.

Lieberman-Aiden, E., Berkum, N.L. van, Williams, L., et al. (2009) Comprehensive mapping of long-range interactions reveals folding principles of the human genome. Science, 326, 289–293.

Li, Z., McKibben, M.T.W., Finch, G.S., Blischak, P.D., Sutherland, B.L. & Barker, M.S. (2021) Patterns and processes of diploidization in land plants. Annual Review of Plant Biology, 72, 387–410.

Manton, I. (1950) The spiral structure of chromosomes. Biological Reviews of the Cambridge Philosophical Society, 25, 486–508.

Marie, D. & Brown, S.C. (1993) A cytometric exercise in plant DNA histograms, with 2C values for 70 species. Biology of the Cell, 78, 41–51.

Michael, T.P. (2014) Plant genome size variation: bloating and purging DNA. Briefings in Functional Genomics, 13, 308–317.

Montgomery, S.A., Tanizawa, Y., Galik, B., et al. (2020) Chromatin organization in early land plants reveals an ancestral association between H3K27me3, transposons, and constitutive heterochromatin. Current Biology, 30, 573–588.e7.

Murray, B.G., Hammett, K.R.W. & Standring, L.S. (1992) Genomic constancy during the development of *Lathyrus odoratus* cultivars. Heredity, 68, 321–327.

Nagl, W. (1985) Chromatin organization and the control of gene activity. International Review of Cytology, 94, 21–56.

Nagl, W. & Bachmann, K. (1980) Condensed chromatin in diploid and allopolyploid microseris species with different genome size: a quantitative electron microscopic study. Theoretical and Applied Genetics, 57, 107–111.

Nagl, W. & Fusenig, H.-P. (1979) Types of chromatin organization in plant nuclei. In W. Nagl, V. Hemleben, and F. Ehrendorfer, eds. Genome and Chromatin: Organization, Evolution, Function: Symposium, Kaiserslautern, October 13–15, 1978. Vienna: Springer Vienna, pp. 221–233.

Nagl, W., Jeanjour, M., Kling, H., Kuhner, S., Michels, I., Muller, T., & Stein, B. (1983) Genome and chromatin organization in higher-plants. Biologisches Zentralblatt, 102, 129–148.

Natarajan, A.T. & Ahnström, G. (1969) Heterochromatin and chromosome aberrations. Chromosoma, 28, 48–61.

Němečková, A., Koláčková, V., Vrána, J., Doležel, J. & Hřibová, E. (2020) DNA replication and chromosome positioning throughout the interphase in three-dimensional space of plant nuclei. Journal of Experimental Botany, 71, 6262–6272.

Novák, P., Neumann, P. & Macas, J. (2020) Global analysis of repetitive DNA from unassembled sequence reads using RepeatExplorer2. Nature Protocols, 15, 3745–3776.

Nowicka, A., Sliwinska, E., Grzebelus, D., Baranski, R., Simon, P.W., Nothnagel, T. & Grzebelus, E. (2016) Nuclear DNA content variation within the genus *Daucus* (Apiaceae) determined by flow cytometry. Scientia Horticulturae, 209, 132–138.

Obermayer, R., Leitch, I.J., Hanson, L. & Bennett, M.D. (2002) Nuclear DNA C-values in 30 species double the familial representation in pteridophytes. Annals of Botany, 90, 209– 217.

Ohri, D., Fritsch, R.M. & Hanelt, P. (1998) Evolution of genome size in *Allium* (Alliaceae). Oesterreichische botanische Zeitschrift, 210, 57–86.

O’Keefe, R.T., Henderson, S.C. & Spector, D.L. (1992) Dynamic organization of DNA replication in mammalian cell nuclei: spatially and temporally defined replication of chromosome-specific alpha-satellite DNA sequences. The Journal of Cell Biology, 116, 1095–1110.

Olszewska, M.J. & Osiecka, R. (1983) The relationship between 2 C DNA content, life cycle type, systematic position and the dynamics of DNA endoreplication in parenchyma nuclei during growth and differentiation of roots in some dicotyledonous herbaceous species. Biochemie und Physiologie der Pflanzen, 178, 581–599.

Orooji, F., Mirzaghaderi, G., Kuo, Y.-T. & Fuchs, J. (2022) Variation in the number and position of rDNA loci contributes to the diversification and speciation in *Nigella* (Ranunculaceae). Frontiers in Plant Science, 13, 917310.

Pellicer, J., Fay, M.F. & Leitch, I.J. (2010) The largest eukaryotic genome of them all? Botanical Journal of the Linnean Society, 164, 10–15.

Pellicer, J. & Leitch, I.J. (2020) The Plant DNA C-values database (release 7.1): an updated online repository of plant genome size data for comparative studies. New Phytologist, 226, 301–305.

Pustahija, F., Brown, S.C., Bogunić, F., et al. (2013) Small genomes dominate in plants growing on serpentine soils in West Balkans, an exhaustive study of 8 habitats covering 308 taxa. Plant Soil, 373, 427–453.

Ramachandran, C. & Narayan, R.K. (1985) Chromosomal DNA variation in *Cucumis*. Theoretical and Applied Genetics, 69, 497–502.

Redi, C.A. & Capanna, E. (2012) Genome size evolution: sizing mammalian genomes. Cytogenetic and Genome Research, 137, 97–112.

Rees, H., Cameron, F.M., Hazarika, M.H. & Jones, G.H. (1966) Nuclear variation between diploid angiosperms. Nature, 211, 828–830.

Ryan Gregory, T. (2011) The Evolution of the Genome, Elsevier.

Sanchez, P.L., Costich, D.E., Friebe, B., Coffelt, T.A., Jenks, M.A. & Gore, M.A. (2014) Genome size variation in guayule and mariola: Fundamental descriptors for polyploid plant taxa. Industrial Crops and Products, 54, 1–5.

Schubert, V., Berr, A. & Meister, A. (2012) Interphase chromatin organisation in *Arabidopsis* nuclei: constraints versus randomness. Chromosoma, 121, 369–387.

Sheval, E.V. (2018) Analysis of сhromosome сondensation/decondensation during mitosis by EdU incorporation in *Nigella damascena* L. seedling roots. Bio-protocol, 8, e2726.

Shinke, N. (1930) On the spiral structure of chromosomes in some higher plants. *Memoirs of the College of Science*, Kyoto Imperial University. Series B, 5, 239–245.

Solovei, I., Kreysing, M., Lanctôt, C., Kösem, S., Peichl, L., Cremer, T., Guck, J. & Joffe, B. (2009) Nuclear architecture of rod photoreceptor cells adapts to vision in mammalian evolution. Cell, 137, 356–368.

Solovei, I., Thanisch, K. & Feodorova, Y. (2016) How to rule the nucleus: divide et impera. Current Opinion in Cell Biology, 40, 47–59.

Sorokin, D.V., Arifulin, E.A., Vassetzky, Y.S. & Sheval, E.V. (2020) Live-cell imaging and analysis of nuclear body mobility. Methods in Molecular Biology, 2175, 1–9.

Sorokin, D.V., Peterlik, I., Tektonidis, M., Rohr, K. & Matula, P. (2018) Non-rigid contour-based registration of cell nuclei in 2-D live cell microscopy images using a dynamic elasticity model. IEEE Transactions on Medical Imaging, 37, 173–184.

Sparvoli, E., Gay, H. & Kaufmann, B.P. (1965) Number and pattern of association of chromonemata in the chromosomes of *Tradescantia*. Chromosoma, 16, 415–435.

Steensel, B. van & Belmont, A.S. (2017) Lamina-associated domains: links with chromosome architecture, heterochromatin, and gene repression. Cell, 169, 780–791.

Temsch, E.M., Greilhuber, J., Hammett, K.R.W. & Murray, B.G. (2008) Genome size in *Dahlia* Cav. (Asteraceae–Coreopsideae). Plant Systematics and Evolution, 276, 157– 166.

Torre, C. de la, Sacristán-Gárate, A. & Navarrete, M.H. (1975) Structural changes in chromatin during interphase. Chromosoma, 51, 183–198.

Van de Peer, Y., Mizrachi, E. & Marchal, K. (2017) The evolutionary significance of polyploidy. Nature Reviews Genetics, 18, 411–424.

Vejdovsky, F. (1912) Zum Problem der Vererbungsträger. Prag: Verlag der Königlich Böhmischen Gesellschaft der Wissenschaften, p. 184.

Weisshart, K., Fuchs, J. & Schubert, V. (2016) Structured illumination microscopy (SIM) and photoactivated localization microscopy (PALM) to analyze the abundance and distribution of RNA polymerase II molecules on flow-sorted *Arabidopsis* nuclei. Bio-protocol, 6, e1725–e1725.

Zhang, X., Pandey, M.K., Wang, J., et al. (2021) Chromatin spatial organization of wild type and mutant peanuts reveals high-resolution genomic architecture and interaction alterations. Genome Biology, 22, 315.

Zonneveld, B.J.M., Leitch, I.J. & Bennett, M.D. (2005) First nuclear DNA amounts in more than 300 angiosperms. Annals of Botany, 96, 229–244.

